# A fluorescent l-2-hydroxyglutarate biosensor

**DOI:** 10.1101/2020.07.07.187567

**Authors:** Zhaoqi Kang, Manman Zhang, Kaiyu Gao, Wen Zhang, Yidong Liu, Dan Xiao, Shiting Guo, Cuiqing Ma, Chao Gao, Ping Xu

## Abstract

l-2-Hydroxyglutarate (l-2-HG) plays important roles in diverse physiological processes, such as carbon starvation response, tumorigenesis, and hypoxic adaptation. Despite its importance and intensively studied metabolism, regulation of l-2-HG metabolism remains poorly understood and a regulator specifically responded to l-2-HG has never been identified. Based on the genomic neighborhood analysis of the gene encoding l-2-HG oxidase (LhgO), LhgR, which represses the transcription of *lhgO*, was identified in *Pseudomonas putida* W619 in this study. LhgR was demonstrated to recognize l-2-HG as its specific effector molecule, and this allosteric transcription factor was then used as a biorecognition element for construction of l-2-HG-sensing FRET sensor. The newly developed l-2-HG sensor can conveniently monitor the concentrations of l-2-HG in various biological samples. In addition to bacterial l-2-HG generation during carbon starvation, biological functions of the l-2-HG dehydrogenase and hypoxia induced l-2-HG accumulation were also revealed by using the l-2-HG sensor in human cells.

## Introduction

l-2-Hydroxyglutarate (l-2-HG) is an important metabolite in various domains of life. In mammals and plants, it is produced by lactate dehydrogenase (LDH) and malate dehydrogenase (MDH)-mediated 2-ketoglutarate (2-KG) reduction under hypoxic conditions^1-5^. In microorganisms, it is a metabolic intermediate of glutarate catabolism produced by a glutarate hydroxylase, CsiD^6-8^. l-2-HG dehydrogenase (L2HGDH) or l-2-HG oxidase (LhgO), an FAD-containing oxidoreductase that converts l-2-HG to 2-KG, plays an indispensable role in the catabolism of l-2-HG^9^. While extensive efforts have been devoted to investigate l-2-HG anabolism and catabolism, the molecular machinery that specifically senses l-2-HG and regulates its metabolism has not been clarified until now.

l-2-HG is an inhibitor of 2-KG dependent dioxygenases with specific pro-oncogenic capabilities^10,11^. Thus, this oncometabolite is viewed as a biomarker for a variety of cancers and its rapid and sensitive measurement in body fluids is of clinical significance^12-15^. Importantly, l-2-HG also has endogenous functions in healthy animal cells. For example, this compound was recently identified to aid in the proliferation and antitumorigenic abilities of CD8^+^ T-lymphocytes^16^, to contribute to relieving the cellular reductive stress^3^, and to coordinate glycolytic flux with epigenetic modifications^17^. Considering the diversity of the roles of l-2-HG in cell metabolism, development and optimization of assays for real-time tracking of this metabolite in living cells is required.

Liquid chromatography-tandem mass spectrometry (LC-MS/MS)^18,19^ and gas chromatography-tandem mass spectrometry (GC-MS/MS)^20,21^ are often used to assess the extracellular concentrations of l-2-HG. These common methods are, at present, time consuming, expensive to perform, and require highly skilled personnel. In addition, these destructive methods are also incompatible with real-time monitoring of the fluctuations of l-2-HG concentrations in intact living cells. In this study, we identified and characterized LhgR, an l-2-HG catabolism regulator in *Pseudomonas putida* W619. Mechanistically, LhgR represses the transcription of LhgO encoding gene *lhgO*. l-2-HG is a specific effector molecule of LhgR and prevents LhgR binding to the promoter region of *lhgO*. Then, we report the development and application of the LhgR based l-2-HG biosensor via Förster resonance energy transfer (FRET), a technology widely applied in the investigation of temporal dynamics of various small molecules, such as potassium^22,23^, glycine^24^, and cAMP^25,26^. The newly developed sensor was used to test l-2-HG concentrations in various biological samples and achieved competitive accuracy and precision. We also took advantage of this biosensor to test predictions about the carbon starvation-induced l-2-HG production in bacteria and to demonstrate hypoxia-induced l-2-HG production by LDH and MDH in human cells. Therefore, the biosensor can also act as a useful tool for real-time measurement of the l-2-HG concentrations in living cells.

## Results

### LhgR regulates l-2-HG catabolism

In this study, bacteria containing LhgO encoding gene *lhgO* were selected to study the regulation of l-2-HG metabolism. Homologs of LhgO can be found in 612 diverse bacterial strains. Similarly organized chromosomal clusters can be found in many bacterial genomes, which contain various combinations of genes related to glutarate metabolism (*csiD, lhgO, gabT, gabD*, and *gabP*) (Fig. 1a). In *Pseudomonas putida* KT2440, the glutarate regulon is regulated by allosteric transcription factor CsiR, which is encoded upstream of *csiD*^27^. The glutarate sensing allosteric transcription factor CsiR and its cognate promoter have been cloned into broad host range vectors to create a glutarate biosensor^28^. Interestingly, a different pattern of *lhgO* gene neighborhood is observed in a few species that do not contain *csiD* homologs (Fig. 1a). For example, a gene encoding a GntR family protein, *lhgR*, is found directly upstream of *lhgO* in *P. putida* W619. The absence of *csiD* gene related to glutarate catabolism made us to anticipate that *lhgO* of *P. putida* W619 might be solely involved in l-2-HG metabolism and be l-2-HG inducible.

**Figure 1.**
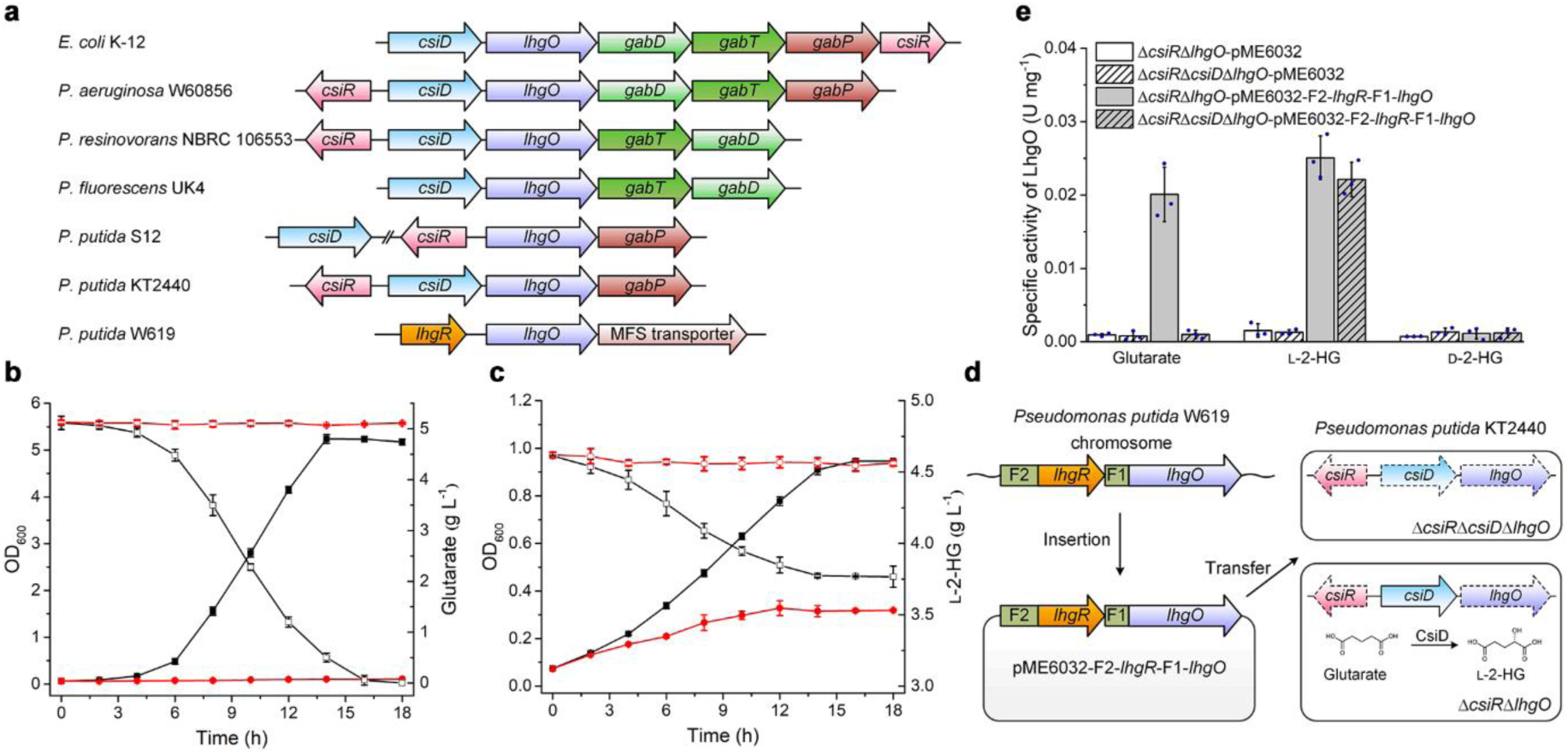
Regulation of l-2-HG catabolism by LhgR in *P. putida* W619. **(a)** Schematic representation of genomic neighborhood analysis of *lhgO* in different bacteria. Orthologs are shown in the same color and the direction of gene transcription is indicated by arrows. CsiR, GntR family allosteric transcription factor regulating glutarate catabolism; CsiD, glutarate hydroxylase; LhgO, l-2-HG oxidase; GabD, succinate semialdehyde dehydrogenase; GabT, 4-aminobutyrate aminotransferase; GabP, 4-aminobutyrate transporter. **(b, c)** Growth of the derivatives of *P. putida* KT2440 in MSMs with glutarate **(b)** or l-2-HG **(c)** as the sole carbon source. Growth (closed symbols) and the consumption of carbon source (open symbols) of *P. putida* KT2440 (Δ*lhgO*) harboring plasmid pME6032-*lhgO* (black lines with squares) and *P. putida* KT2440 (Δ*lhgO*) harboring empty plasmid pME6032 (red lines with circles) were measured in MSMs containing 5 g L^-1^ glutarate **(b)** or l-2-HG **(c)** as the sole carbon source. **(d)** Schematic representation of the construction of pME6032-F2-*lhgR*-F1-*lhgO* and its transfer into *P. putida* KT2440 (Δ*csiR*Δ*lhgO*) and *P. putida* KT2440 (Δ*csiR*Δ*csiD*Δ*lhgO*) by electroporation. The deleted genes in *P. putida* KT2440 are indicated by dashed lines. The reaction catalyzed by CsiD is also demonstrated. **(e)** The activities of LhgO in *P. putida* KT2440 (Δ*csiR*Δ*lhgO*) and *P. putida* KT2440 (Δ*csiR*Δ*csiD*Δ*lhgO*) harboring either plasmid pME6032-F2-*lhgR*-F1-*lhgO* or empty plasmid pME6032 grown in MSM with glutarate, l-2-HG, or D-2-HG as the sole carbon source. All data shown are means ± standard deviations (s.d.) (n = 3 independent experiments).

The *lhgO* gene in *P. putida* W619 was cloned into pME6032 vector, and the resulting plasmid was transferred into *P. putida* KT2440 (Δ*lhgO*). As shown in Fig. 1b-c, complement of *lhgO* in *P. putida* W619 could restore glutarate and l-2-HG utilization abilities of *P. putida* KT2440 (Δ*lhgO*), confirming that *lhgO* encodes a functional l-2-HG catabolic enzyme. To identify the function of LhgR in *P. putida* W619, the gene segment F2-*lhgR*-F1-*lhgO*, which contains the promoter of *lhgR* (F2), *lhgR*, the promoter of *lhgO* (F1), and *lhgO*, was cloned into pME6032 vector, and the resulting plasmid was transferred into different derivatives of *P. putida* KT2440 (Fig. 1d). As shown in Fig. 1e, exogenous l-2-HG, but not its mirror-image enantiomer D-2-HG, can induce the expression of *lhgO* in the gene segment F2-*lhgR*-F1-*lhgO* and restore LhgO activity in *P. putida* KT2440 (Δ*csiR*Δ*lhgO*). In addition, the activity of LhgO was also detected in *P. putida* KT2440 (Δ*csiR*Δ*lhgO*) harboring pME6032-F2-*lhgR*-F1-*lhgO* when cultured with glutarate as the sole carbon source. However, no activity of LhgO was detected in *P. putida* KT2440 (Δ*csiR*Δ*csiD*Δ*lhgO*), in which the key gene responsible for l-2-HG production from glutarate was deleted. These results indicated that LhgR represses the expression of LhgO and l-2-HG, but nor D-2-HG or glutarate, might be the effector molecule of LhgR.

### LhgR specifically responds to l-2-HG

To determine whether LhgR directly interacts with the promoter region of *lhgO*, LhgR in *P. putida* W619 was overexpressed in *E. coli* BL21(DE3) and purified by Ni-chelating chromatography (Fig. 2a). Based on the results of gel filtration and sodium dodecyl sulfate-polyacrylamide gel electrophoresis (SDS-PAGE), LhgR behaved as a dimer (Fig. 2b). Subsequently, electrophoretic mobility shift assays (EMSAs) were conducted using *lhgO* promoter (F1) and purified LhgR. As shown in Fig. 2c, LhgR bound to F1 in a concentration-dependent manner. LhgR completely shifted fragment F1 when an 8-fold molar excess was used. A DNase I footprinting assay was also performed using purified LhgR and fragment F1. A protected region containing palindromic N_yGTNx>ACNy_ consensus binding motif of GntR-family allosteric transcription factor^29^, 5^’^-TA**GT**CTG**AC**AA-3’, was observed (Fig. 2d). In addition, LhgR also bound to its promoter (F2) in EMSAs and a similar consensus binding motif, 5^’^-TT**GT**CTG**AC**AA-3’, was protected in DNase I footprinting assay (Supplementary Fig. 1a-b).

**Figure 2.**
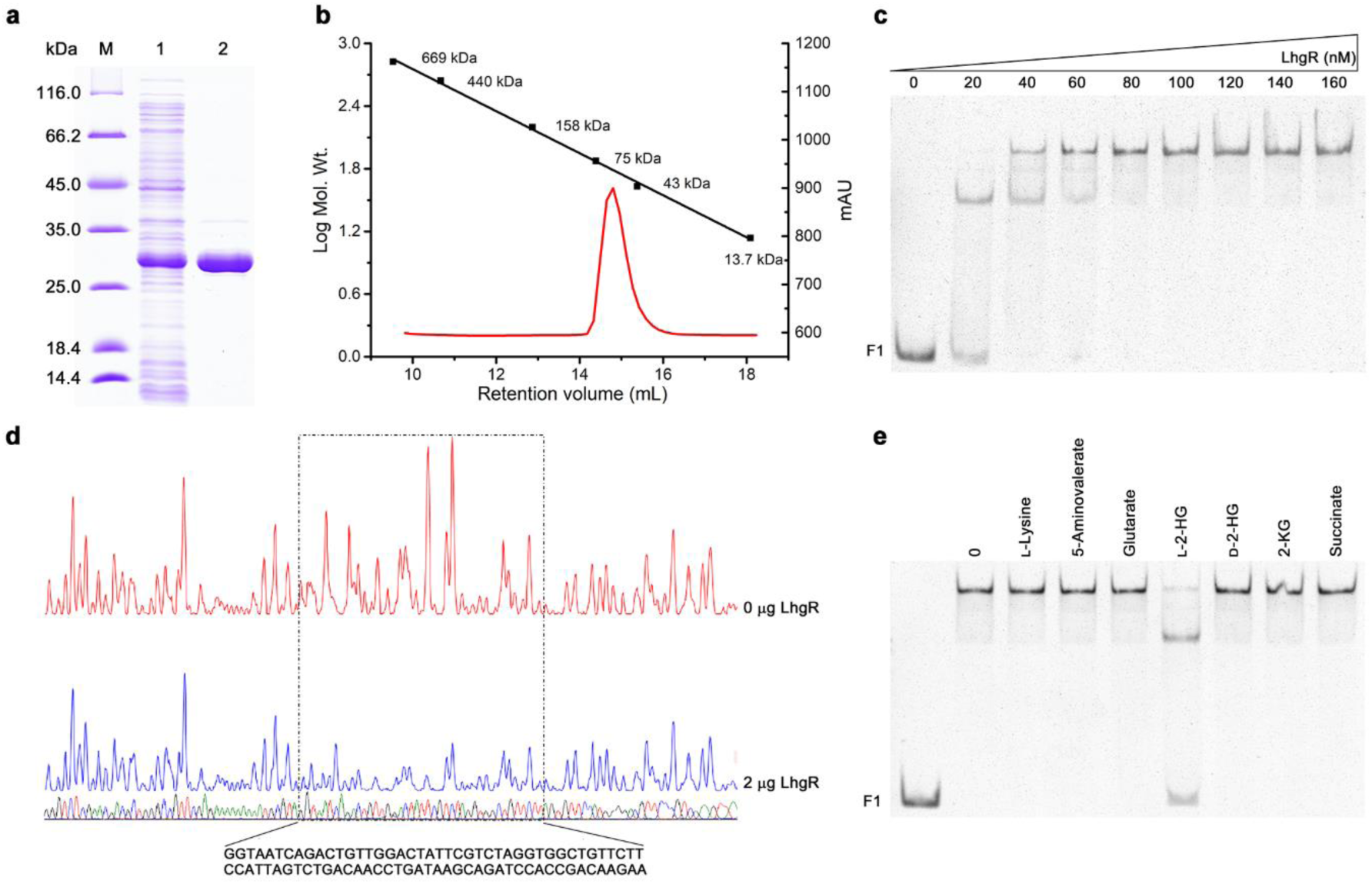
Purification and characterization of LhgR. **(a)** SDS-PAGE analysis of the purification of LhgR. Lane M, molecular weight markers; lane 1, crude extract of *E. coli* BL21(DE3) harboring pETDuet-*lhgR*; lane 2, purified His_6_-tagged LhgR using a HisTrap column. **(b)** Gel-filtration chromatography of the purified LhgR with the Superdex 200 10/300 GL column. Red curve, chromatogram of purified LhgR; Black line, standard curve for protein molecular mass standards. **(c)** LhgR can bind to the *lhgO* promoter region. F1 fragment containing the *lhgO* promoter region (10 nM) was titrated by purified LhgR (0, 20, 40, 60, 80, 100, 120, 140, 160 nM). **(d)** DNase I footprinting analysis of LhgR binding to the *lhgO* promoter region. The F1 fragment was labeled with 6-carboxyfluorescein (FAM) and incubated with 2 μg LhgR (blue line) or without LhgR (red line). The region protected by LhgR is indicated with a dotted box. **(e)** l-2-HG prevent LhgR binding to the *lhgO* promoter region. EMSAs were carried out with F1 fragment (10 nM) and purified LhgR (60 nM) in the absence of any other tested compounds (0) and in the presence of 50 mM different compounds. The leftmost lane without LhgR was used as the control.

The effects of l-2-HG, D-2-HG, glutarate, 2-KG, L-lysine, 5-aminovalerate, and succinate on LhgR binding to the *lhgO* promoter region F1 were also assessed by EMSAs. The release of LhgR from fragments F1 was observed only in the presence of l-2-HG (Fig. 2e). These results indicated that l-2-HG can specifically prevent the binding of LhgR to the promoter of *lhgO* and induce its expression. In particular, LhgR might self-repress its expression and l-2-HG can also contribute in inducing the expression of *lhgR* (Supplementary Fig. 1c).

### Design and optimization of the l-2-HG-sensing reporter

FRET sensors, which combine a ligand-binding moiety and a pair of donor–acceptor fluorescent pair, allow measurement of ligand concentrations based on the ligand-binding induced changes of FRET efficiency^22-26^. In this study, the l-2-HG-sensing fluorescent reporter (LHGFR) was constructed by fusion of the optimized cyan and yellow fluorescent protein variants, mTFP^30^ and Venus^31^, to the N- and C-terminus of LhgR (Supplementary Fig. 2). This first LHGFR was named as LHGFR_0N0C_, where the subscript indicates the number of amino acids truncated from the N- and C-terminus of LhgR. Subsequently, LHGFR_0N0C_ was overexpressed in *E. coli* BL21(DE3) and purified by a Ni-chelating chromatographic column (Supplementary Fig. 3). The addition of l-2-HG could reduce the emission peak at 492 nm of mTFP and increase the emission peak at 526 nm of Venus (Supplementary Fig. 4), indicating that l-2-HG binding to LHGFR_0N0C_ increases FRET between the fluorophores (Fig. 3a). In addition, l-2-HG increased the emission ratio of Venus to mTFP in a dose-dependent manner, with a maximum ratio change (Δ*R*_*max*_) of 11.47 ± 0.38%, an apparent dissociation constant (*K*_*d*_) of 2.74 ± 0.73 μM, and a Hill slope close to 1 (Fig. 3b).

**Figure 3.**
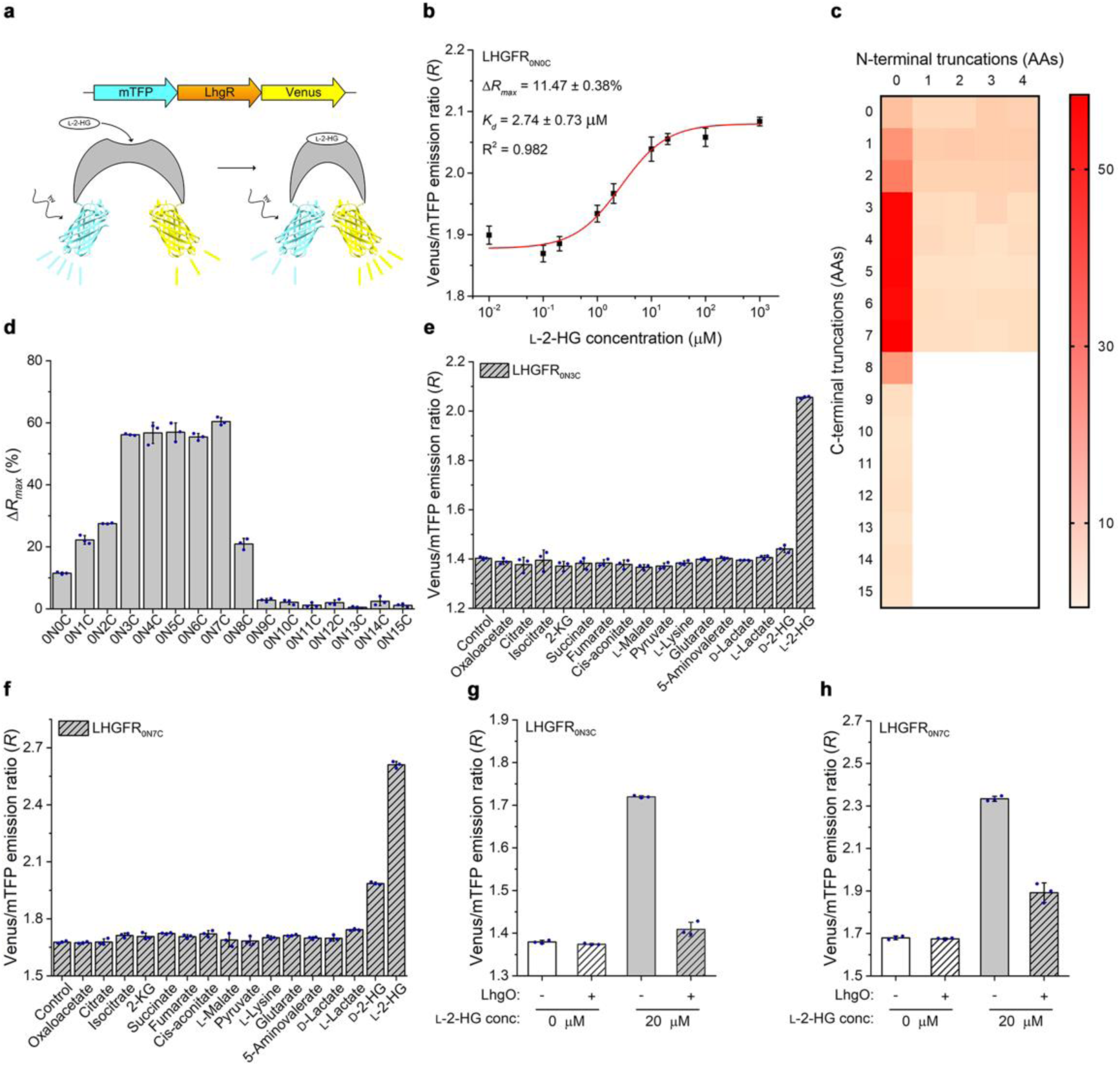
Design and optimization of the LHGFR. **(a)** Schematic representation of the predicted conformational change of FRET-based l-2-HG biosensor LHGFR in the absence or presence of l-2-HG. **(b)** Dose-response curve of purified LHGFR_0N0C_ for increasing concentrations (10 nM to 1 mM) of l-2-HG in 50 mM Tris-HCl buffer (pH 7.4). The emission ratio of Venus to mTFP increased (430 nm excitation) after l-2-HG binding. **(c)** Heap map of the truncating the N-and C-terminal amino acids of LhgR to Δ*R*_*max*_. Color indicates the value of Δ*R*_*max*_ and white indicates the untested variants. **(d)** Comparison of the Δ*R*_*max*_ of a set of l-2-HG biosensor variants based on the C-terminal amino acid truncated of LhgR. **(e, f)** Evaluation of the specificity of the purified LHGFR_0N3C_ **(e)** and LHGFR_0N7C_ **(f)**. The emission ratio changes of both biosensors were measured in the presence of 240 μM of D-lactate, L-lactate, D-2-HG, or different intermediates of TCA cycle and L-lysine catabolism, respectively. The rightmost column in the presence of l-2-HG was used as the control. **(g, h)** Reversal of l-2-HG binding with LHGFR by conversion of l-2-HG to 2-KG. The emission ratio of purified LHGFR_0N3C_ **(g)** and LHGFR_0N7C_ **(h)** was recorded in the absence and presence of 20 μM l-2-HG before and after the addition of 5 μM purified LhgO for 10 min. All data shown are means ± s.d. (n = 3 independent experiments).

To increase the magnitude of responses, LHGFR was optimized by truncating N- and C-terminal amino acids of LhgR or by adding a series of artificial linkers between LhgR and various fluorescent proteins^23,32-34^ (Fig. 3c and Supplementary Fig. 5). Truncation of just three to seven C-terminal amino acids in LhgR could significantly increase Δ*R*_*max*_ of the sensor (Fig. 3c-d and Supplementary Fig. 5). Among the five sensors with increased response magnitude values, LHGFR_0N7C_ was the best mutant with a higher Δ*R*_*max*_ of 60.37 ± 1.30% (Fig. 3d and Supplementary Fig. 6a-e). In addition, LHGFR_0N3C_ was also a promising sensor with a high Δ*R*_*max*_ of 56.13 ± 0.29% and a high *K*_*d*_ of 29.33 ± 1.24 μM (Supplementary Fig. 6a). The unique *K*_*d*_ of LHGFR_0N3C_ might make this sensor usable in monitoring l-2-HG at a high dynamic range.

Then, a set of intermediates of TCA cycle and L-lysine catabolism were used to examine the specificity of LHGFR_0N3C_ and LHGFR_0N7C_. None of oxaloacetate, citrate, isocitrate, 2-KG, succinate, fumarate, cis-aconitate, L-malate, pyruvate, L-lysine, 5-aminovalerate, glutarate, D-lactate, and L-lactate induced emission ratio changes in LHGFR_0N3C_ or LHGFR_0N7C_ (Fig. 3e-f). Although D-2-HG elicited small changes in the emission ratio of LHGFR_0N7C_ at a concentration of 240 μM, dose-response curves of LHGFR_0N3C_ and LHGFR_0N7C_ for D-2-HG revealed insignificant binding of D-2-HG and *K*_*d*_ values much higher than that for l-2-HG (Supplementary Fig. 7a-b). l-2-HG-dependent emission ratio changes of LHGFR_0N3C_ or LHGFR_0N7C_ were significantly reversible by l-2-HG oxidation catalyzed by 5 μM LhgO (Fig. 3g-h) and both biosensors were stable for the detection of l-2-HG from pH 6.0 to 8.0 (Supplementary Fig. 8a-b). The high specificity, reversibility, and stability of LHGFR_0N3C_ and LHGFR_0N7C_ indicated the availability of these biosensors in monitoring dynamic changes in l-2-HG under physiological conditions.

### Characterization of LHGFR in biological samples

Next, we investigated whether LHGFR could be used to quantify l-2-HG concentrations in different biological samples. When l-2-HG with increasing concentrations (0 to 2 mM) were added into serum and urine samples of healthy adults, the response curves were nearly identical with that in assay buffer for both LHGFR_0N3C_ and LHGFR_0N7C_ (Fig. 4a-d, Supplementary Fig. 6a and Fig. 6e). Thus, quantitative determination of l-2-HG could be conducted by mixing the target samples with LHGFR and measuring the emission ratios with a conventional fluorescence microplate reader. Based on the response curves established for l-2-HG quantification, both biosensors were used to assay the concentrations of l-2-HG in human serum and urine (Fig. 4e-f). The results of LHGFR_0N3C_ and LHGFR_0N7C_ showed close agreement with the results of LC-MS/MS, the current standard method for clinical assays of l-2-HG. Besides its accuracy and precision in l-2-HG quantification (Supplementary Table 1), LHGFR_0N7C_ also exhibited superior applicability in its lower limits of detection (LOD) (Supplementary Table 2).

**Figure 4.**
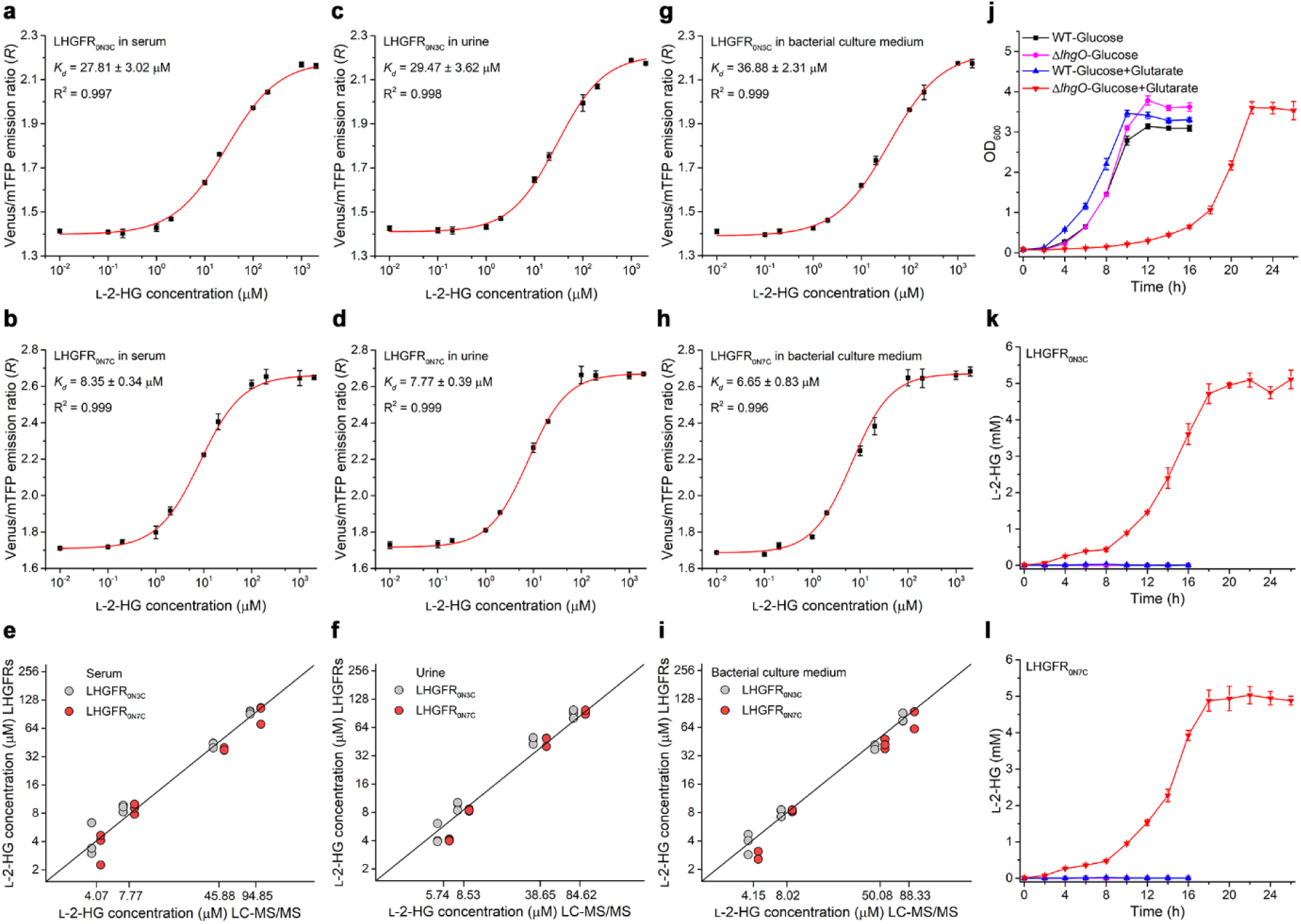
Validation of purified LHGFR for determination l-2-HG levels in body fluids and bacterial culture system. **(a-d)** Dose-response curves of purified LHGFR_0N3C_ and LHGFR_0N7C_ for increasing concentrations (10 nM to 2 mM) of l-2-HG in serum **(a, b)** and urine **(c, d). (e, f)** Comparison between the quantitative results of l-2-HG in serum **(e)** and urine **(f)** by LC-MS/MS and LHGFR. The gray circles and red circles represent the quantitative results of LHGFR_0N3C_ and LHGFR_0N7C_, respectively. Black line indicates a reference line. **(g, h)** Dose-response curves of purified LHGFR_0N3C_ **(g)** and LHGFR_0N7C_ **(h)** for increasing concentrations (10 nM to 2 mM) of l-2-HG in bacterial culture medium. **(i)** Comparison between the quantitative results of l-2-HG in bacterial culture medium by LC-MS/MS and LHGFR. **(j)** Growth of *P. putida* KT2440 and its *lhgO* mutant in medium containing 20 mM glucose and 5 mM glutarate as the carbon sources. **(k, l)** Determination of extracellular l-2-HG accumulation of *P. putida* KT2440 and its *lhgO* mutant by purified LHGFR_0N3C_ **(k)** and LHGFR_0N7C_ **(l)**. All data shown are means ± s.d. (n = 3 independent experiments).

**Figure 5.**
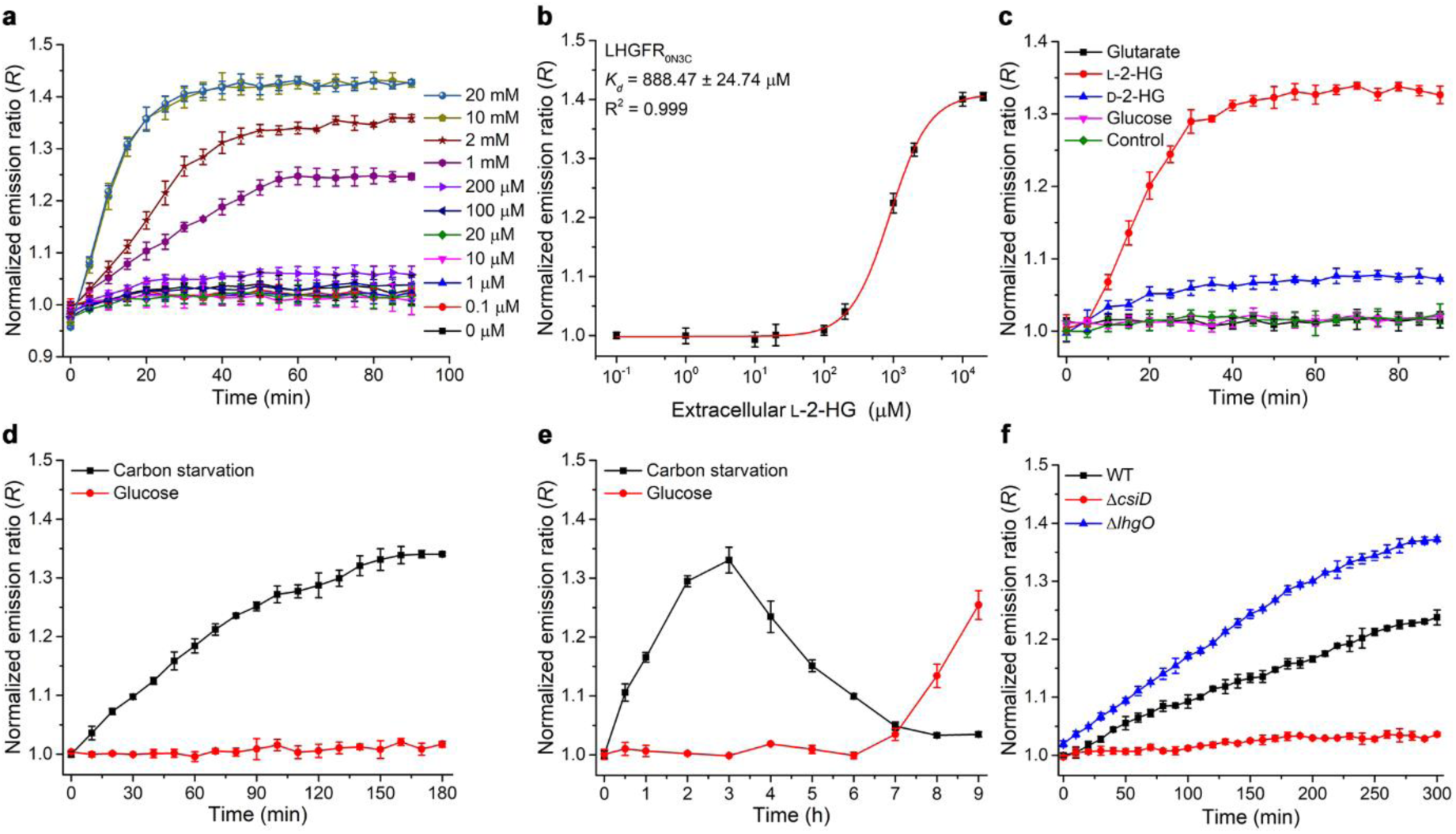
Monitoring l-2-HG fluctuations in living bacteria by LHGFR. **(a)** Time course of the emission ratio changes of LHGFR_0N3C_ expressed in *E. coli* BL21(DE3) in response to exogenous l-2-HG addition. All ratios were normalized to the control (ratio in the absence of l-2-HG at time point zero). **(b)** Normalized dose-response curve of LHGFR_0N3C_ expressed in *E. coli* BL21(DE3) for increasing concentrations (100 nM to 20 mM) of l-2-HG at time point 60 min. **(c)** Time course of the emission ratio changes of LHGFR_0N3C_ expressed in *E. coli* BL21(DE3) in response to the addition of 1 mM glutarate, l-2-HG, D-2-HG, or glucose. All data were normalized to the control (ratio in the absence of any tested compounds at time point zero). **(d)** Detection of carbon starvation-induced l-2-HG accumulation over time by LHGFR_0N3C_ expressed in *E. coli* BL21(DE3). Emission ratio changes of LHGFR_0N3C_ were measured when cultured in carbon starvation medium (black line) and medium with 20 mM glucose (red line). All data were normalized to samples under carbon starvation condition at time point zero. **(e)** Long-term detection of l-2-HG fluctuations by LHGFR_0N3C_ expressed in *E. coli* BL21(DE3). All data were normalized to samples under carbon starvation condition at time point zero. **(f)** Identification of the roles of CsiD and LhgO in endogenous l-2-HG catabolism during carbon starvation by LHGFR_0N3C_. Emission ratio changes of LHGFR_0N3C_ expressed in *E. coli* MG1655(DE3) wild-type (black line), *E. coli* MG1655(DE3) (Δ*csiD*) (red line), and *E. coli* MG1655(DE3) (Δ*lhgO*) (blue line) were measured in carbon starvation medium. All data were normalized to time point zero of wild-type strain and shown as means ± s.d. (n = 3 independent experiments).

**Figure 6.**
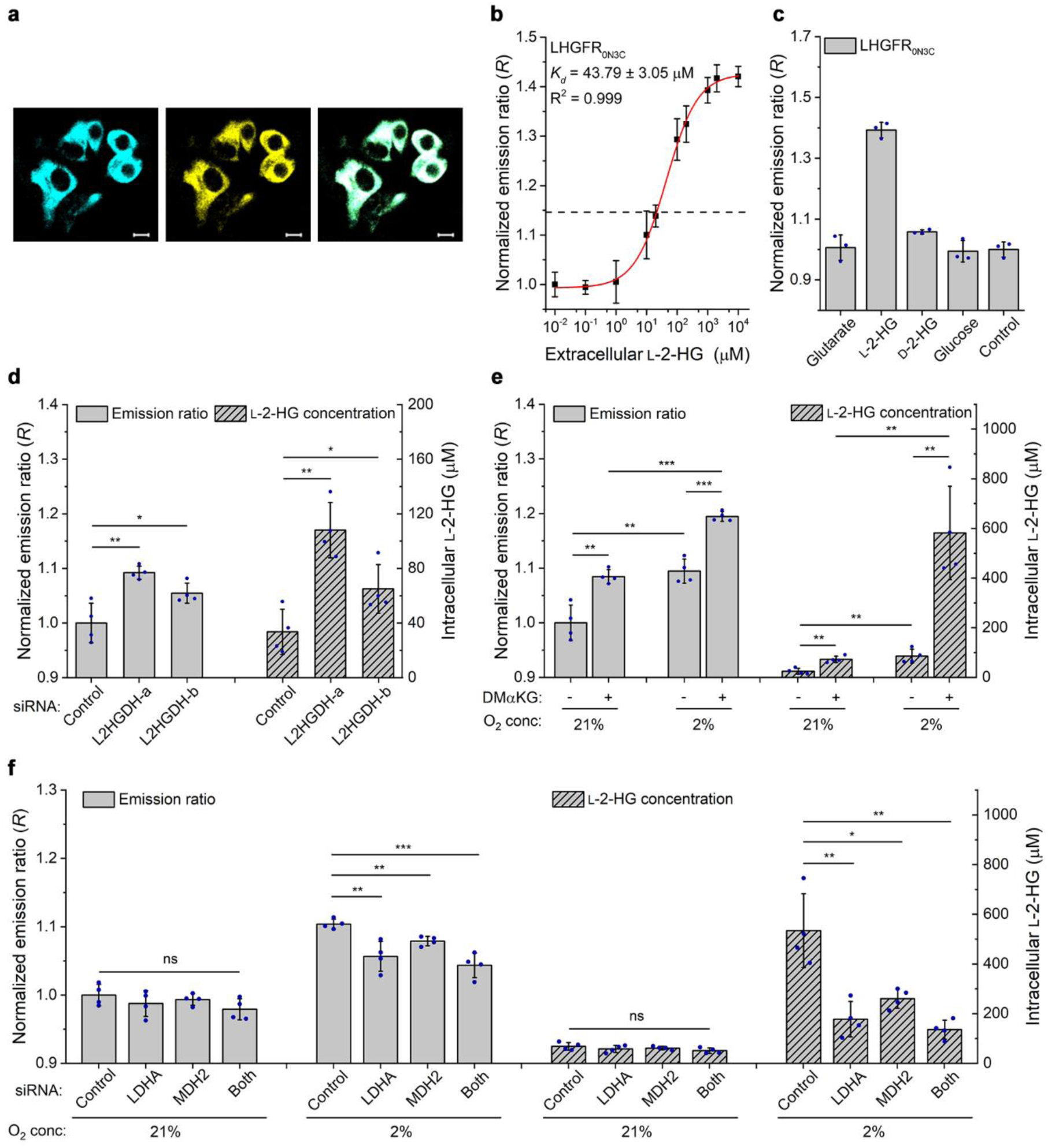
Monitoring l-2-HG fluctuations in human cells by LHGFR. **(a)** Confocal microscopy images of LHGFR_0N3C_-expressing HEK293FT cells. The images are represented as mTFP channel, Venus channel, and overlay channel from left to right. Scale bar, 10 μm. **(b)** Normalized dose-response curve of LHGFR_0N3C_ expressed in HEK293FT cells with increasing concentrations (10 nM to 10 mM) of l-2-HG. Cells were permeabilized with 10 μM digitonin. The emission ratio of non-permeabilized HEK293FT cells under physiological conditions is indicated with black dash line. **(c)** Responses of LHGFR_0N3C_ expressed in HEK293FT cells to exogenously added 1 mM glutarate, l-2-HG, D-2-HG, and glucose. All data were normalized to the control (ratio in the absence of any tested compounds). **(d)** Identification of the function of L2HGDH in l-2-HG catabolism by LHGFR_0N3C_. The emission ratio was measured after co-transfecting siRNA targeting L2HGDH and LHGFR_0N3C_ for 48 h. **(e)** Detection of hypoxia-induced l-2-HG accumulation by LHGFR_0N3C_. Emission ratio changes were recorded after LHGFR_0N3C_-expressing HEK293FT cells cultured in normoxia or hypoxia in the absence and presence of 5 mM dimethyl-2-ketoglutarate (DMαKG) for 24 h. Emission ratio was normalized to normoxic condition without DMαKG. **(f)** Identification of the functions of LDHA and MDH2 in l-2-HG anabolism by LHGFR_0N3C_. HEK293FT cells were cultured in the presence of 5 mM DMαKG. Emission ratio was normalized to normoxic condition treated with negative siRNA. All data shown are means ± s.d. (n = 3, 3, 4, 4, and 4 independent experiments for **b, c, d, e**, and **f**). *, *P* < 0.05 in two-tailed *t* test; **, *P* < 0.01 in two-tailed *t* test; ***, *P* < 0.001 in two-tailed *t* test; ns, no significant difference (*P* ≥ 0.05 in two-tailed *t* test).

In a previous report, l-2-HG was confirmed to be a metabolic intermediate of glutarate metabolism in *P. putida* KT2440^7^. LHGFR_0N3C_ and LHGFR_0N7C_ also exhibited high accuracy and precision in quantification of l-2-HG in bacterial culture medium (Fig. 4g-i). When cultured in medium containing 20 mM glucose and 5 mM glutarate as carbon sources, the growth of *P. putida* KT2440 (Δ*lhgO*) was significantly delayed which might be due to the possible toxicity of accumulated l-2-HG (Fig. 4j). Nearly identical results of l-2-HG quantification were also obtained by either using LHGFR_0N3C_ or LHGFR_0N7C_ (Fig. 4k-l). Mutual corroboration between the two biosensors further confirmed their applicabilities in *in vitro* l-2-HG quantification of various biological samples.

### Monitoring l-2-HG fluctuations in living bacteria by LHGFR

We investigated whether the FRET-based l-2-HG sensor LHGFR could detect possible variations of l-2-HG in living bacteria. The biosensor LHGFR_0N3C_ was expressed in *E. coli* BL21(DE3). Exogenous l-2-HG was added to the culture system of *E. coli* BL21(DE3) to achieve concentrations between 0 and 20 mM and the emission ratio was continuously recorded. As shown in Fig. 5a, exogenous l-2-HG at concentrations lower than 10 mM could increase emission ratio in a dose-dependent manner. The apparent *K*_*d*_ of LHGFR_0N3C_ expressed in *E. coli* BL21(DE3) was determined to be 888.47 ± 24.74 μM by fitting emission ratios against exogenous l-2-HG concentrations (Fig. 5b). Thus, the intracellular l-2-HG concentrations might be approximately 30-fold lower than the external supply, assuming that the affinity of LHGFR is not affected in the cytosolic environment^35^. The specificity of LHGFR_0N3C_ expressed in *E. coli* BL21(DE3) was also characterized. As shown in Fig. 5c, only exogenous l-2-HG could significantly increase the emission ratio in *E. coli* BL21(DE3), while glutarate, D-2-HG, and glucose could not.

Besides being a metabolic intermediate of exogenous glutarate catabolism in *P. putida* KT2440, l-2-HG is also reported as a metabolite produced from endogenous L-lysine during carbon starvation of *E. coli*^36,37^. Thus, whether carbon starvation could induce intracellular l-2-HG accumulation of *E. coli* was investigated. As shown in Fig. 5d, no change in the emission ratio was detected when 20 mM glucose was added to the culture system. However, the emission ratio increased during carbon starvation of *E. coli* BL21(DE3), suggesting that carbon starvation induced l-2-HG production. The emission ratios also increased after culturing *E. coli* cells for 6 h with glucose addition (Fig. 5e), which might be due to carbon starvation induced by depletion of exogenous glucose. In addition, the emission ratio increased at the beginning of carbon starvation, reaching a maximum value at 3 h and then decreased to initial levels at 8 h (Fig. 5e). These results confirmed that l-2-HG is a temporary metabolite during carbon starvation and LHGFR_0N3C_ can indeed detect a change in intracellular l-2-HG concentrations in real time.

To identify whether carbon starvation induced endogenous l-2-HG production also results from the glutarate hydroxylase activity of CsiD, gene *csiD* was disrupted and LHGFR_0N3C_ was expressed in *E. coli* MG1655(DE3). Gene *lhgO* in *E. coli* MG1655(DE3) was also disrupted to investigate its role in endogenous l-2-HG catabolism. As expected, the emission ratio of LHGFR_0N3C_ in *E. coli* MG1655(DE3) (Δ*csiD*) remained unaffected during carbon starvation, whereas disruption of *lhgO* significantly increased the emission ratio of LHGFR_0N3C_ in *E. coli* MG1655(DE3) (Δ*lhgO*) (Fig. 5f), confirming the roles of CsiD and LhgO in endogenous l-2-HG catabolism during carbon starvation. The performance of LHGFR_0N7C_ in monitoring l-2-HG fluctuations in living bacteria was also studied and similar results were acquired (Supplementary Fig. 9).

### Monitoring l-2-HG production in human cells by LHGFR

Next, both the biosensors were expressed in HEK293FT cells to monitor l-2-HG levels inside living human cells. As shown in Fig. 6a and Supplementary Fig. 10a, both biosensors were successfully expressed in HEK293FT cells, mainly localized in the cytosol. LHGFR_0N3C_ and LHGFR_0N7C_ were calibrated by adding increasing concentrations (0 to 10 mM) of l-2-HG to 10 μM digitonin-permeabilized HEK293FT cells. The apparent *K*_*d*_ of LHGFR_0N3C_ and LHGFR_0N7C_ expressed in HEK293FT cells were 43.79 ± 3.05 μM and 9.35 ± 0.67 μM, respectively (Fig. 6b and Supplementary Fig. 10b). Based on the emission ratio of non-permeabilized HEK293FT cells under physiological conditions, the basal l-2-HG concentration was determined to be 22.95 ± 11.22 μM in LHGFR_0N3C_-expressing cells (Fig. 6b), and 20.24 ± 8.39 μM in LHGFR_0N7C_-expressing cells (Supplementary Fig. 10b). As shown in Fig. 6c and Supplementary Fig. 10c, only exogenous l-2-HG could significantly increase the emission ratio in 10 μM digitonin-permeabilized HEK293FT, further suggesting the high specificity of these two biosensors.

LHGFR_0N3C_ and LHGFR_0N7C_ were then used to detect l-2-HG fluctuations associated with the loss-of-function of L2HGDH, the only reported enzyme which can catabolize l-2-HG in human cells. HEK293FT cells were co-transfected with siRNA targeting L2HGDH and LHGFR_0N3C_ or LHGFR_0N7C_. As shown in Fig. 6d and Supplementary Fig. 10d, the transfection of siRNA targeting L2HGDH increased the emission ratio of both the biosensors, indicating an accumulation of intracellular l-2-HG due to a decrease in L2HGDH levels (Fig. 6d and Supplementary Fig. 10d).

The sensitivity of LHGFR to changes in hypoxia-induced production of l-2-HG was also studied. LHGFR_0N3C_, which has a wide detection range due to its high *K*_*d*_, was used for this purpose. The emission ratio of LHGFR_0N3C_ in HEK293FT cells after 24 h exposure to 2% oxygen was higher than the ratio obtained under normoxic conditions and hypoxia induced a 3.5-fold increase of l-2-HG (Fig. 6e). In addition, exogenous cell-permeable dimethyl-2-ketoglutarate significantly increased the emission ratio of LHGFR_0N3C_ under hypoxic conditions, suggesting that hypoxia-induced l-2-HG might originate from 2-KG (Fig. 6e).

LDHA and MDH2 have been reported to participate in hypoxia-induced l-2-HG production due to their “promiscuous” catalytic activities^1^. In support of this conclusion, siRNAs targeting LDHA and MDH2 were transfected separately or in combination into LHGFR_0N3C_-expressing HEK293FT cells. As shown in Fig. 6f, decrease of LDHA and MDH2 significantly reduced the emission ratio of LHGFR_0N3C_ under hypoxic conditions, suggesting that these enzymes indeed contribute to the production of l-2-HG from 2-KG.

## Discussion

Bacteria have evolved to respond and catabolize a wide range of metabolites. The availability of genomic information from different organisms provided researchers with a new route to develop robust predictions for new transcriptional regulators and their physiological functions. Here, we used a genomic neighborhood analysis approach combined with genetic and biochemical techniques to uncover a novel transcriptional regulator of l-2-HG catabolism. The identified transcriptional regulator, LhgR, was present directly upstream of LhgO in *P. putida* W619 (Fig. 1a). It specifically binds as a dimer to the promoter region of *lhgO* and represses the transcription of *lhgO* gene. l-2-HG interferes with the DNA-binding activity of LhgR and induces expression of LhgO. To our knowledge, LhgR is the first known example of an allosteric transcription factor which specially responses to l-2-HG in all domains of life. This finding showed the application of a collection of sequenced genomes in identification of novel transcriptional regulators and the approach can be expanded to target other transcriptional regulators in diverse bacteria.

l-2-HG is a harbinger of altered metabolism and participates in the pathogenesis of l-2-hydroxyglutaric acidurias and cancer^12-15,38^. Standard methods to measure l-2-HG are based on MS-based techniques which are time consuming and requires expensive highly skilled workers. LhgR can bind to l-2-HG and then undergo conformational changes, which in turn affects DNA-binding. Thus, a FRET biosensor, LHGFR_0N0C_, utilizing the allosteric transcription factor LhgR as an l-2-HG biorecognition element was constructed for a convenient assay of l-2-HG concentrations. As a ratiometric sensor, the emission ratio changes of LHGFR is not affected by the amount of sensor in biological samples or in living cells, and thus allows more accurate measurements. The biosensor was optimized by truncating the N- and C-terminal domains of LhgR or by adding an artificial linker to the N- and C-terminal regions of LhgR. The optimized variants, LHGFR_0N3C_ and LHGFR_0N7C_, increased Δ*R*_*max*_ from 11.47 ± 0.38% to 56.13 ± 0.29% and 60.37 ± 1.30%, respectively, significantly improving the sensitivity for l-2-HG detection (Fig. 3b, Supplementary Fig. 6a and Fig. 6e). Besides signal recognition, signal transduction is also an essential aspect for the development of biosensors. Various biosensing systems, including CRISPR-Cas12a- and allosteric transcription factors-mediated small molecule detector^39^, allosteric transcription factors-based nicked DNA-template-assisted signal transduction^40^, and quantum-dot-allosteric transcription factors-FRET^41^, have been reported. Other biosensors based on l-2-HG responding to LhgR and the corresponding transduction mechanisms could also be developed for detection of l-2-HG.

As an initial test of LHGFR for quantitative analysis of l-2-HG, the responses of LHGFR_0N3C_ and LHGFR_0N7C_ to varying concentrations of l-2-HG in artificial urine and serum (prepared through addition of l-2-HG to the serum and urine of healthy adults) were measured. Our data indicated that the detection ranges of LHGFR_0N3C_ and LHGFR_0N7C_ for serum and urine were 5.84-4000 μM and 15.74-4000 μM, 1.68-400 μM and 0.92-400 μM, respectively (Supplementary Table 2). The reported concentration of l-2-HG in plasma of patients with l-2-hydroxyglutaric aciduria and l-2-HG-associated brain malignancies is about 7-84 μM^38,42^. Thus, l-2-HG quantification using the highly sensitive LHGFR, especially LHGFR_0N7C_, requires simply mixing traces of serum with LHGFR and then detecting changes in ratios using a fluorescence microplate reader. Besides higher detection sensitivity, LHGFR also has superior accuracy and precision over previous MS-based methods for l-2-HG detection (Fig. 4 and Supplementary Table 1). Being genetically-encoded, l-2-HG sensors can be produced in great quantities by recombinant bacteria with low cost and could be applied in future rapid and sensitive clinical diagnosis of l-2-HG-related diseases.

Then, LHGFR_0N3C_ and LHGFR_0N7C_ were expressed in HEK293FT and digitonin was used to induce cell permeabilization and deplete intracellular l-2-HG for *in vivo* response curves construction. The calibrated *in vivo* response curves of both biosensors in digitonin-permeabilized HEK293FT cells gave similar *K*_*d*_ of those in *in vitro* assays. Based on the *in vivo* response curves of LHGFR_0N3C_ or LHGFR_0N7C_, the basal l-2-HG concentration in HEK293FT cells under physiological conditions was 22.95 ± 11.22 μM or 20.24 ± 8.39 μM, respectively (Fig. 6b and Supplementary Fig. 10b). l-2-HG quantification in living HEK293FT cells using LHGFR got similar results to that acquired using MS-based approaches after complicated sample handling and data analysis^2,3^. Considering the fact that hypoxic cells, activated T-cells, and certain cancer cells might accumulate l-2-HG to high concentrations^2-4,12-14,16,43^, LHGFR_0N3C_ with high *K*_*d*_ and detection range might be a more viable alternative for the *in vivo* assay of l-2-HG concentrations.

The potential of LHGFR for real-time monitoring of fluctuations in intracellular l-2-HG concentrations was illustrated by using bacterial cells and HEK293FT cells. It was revealed that carbon starvation also induced temporary intracellular accumulation of l-2-HG in *E. coli* cells. CsiD and LhgO played indispensable roles in endogenous anabolism and catabolism of l-2-HG, respectively (Fig. 5f and Supplementary Fig. 9f). In addition, it was identified that growth of the strain containing *lhgO* mutation was inhibited when high levels of l-2-HG was present (Fig. 4j-l). Besides being a pathogenic metabolite inducing various cancers and l-2-hydroxyglutaric aciduria in humans^12-15,38,42^, l-2-HG would also be a toxic metabolite to bacterial cells. l-2-HG catabolizing enzymes, including L2HGDH in humans^9^, dL2HGDH in *Drosophila*^17^, and LhgO in *P. putida*^7,8^ and *E. coli*^6,44^, might all exist as detoxification proteins of l-2-HG. The functions of L2HGDH in l-2-HG catabolism and LDHA and MDH2-mediated 2-KG reduction in hypoxia-induced l-2-HG production were also confirmed in HEK293FT cells using LHGFR as an indicator of l-2-HG. LHGFR can be added in the emerging list of metabolite sensors that have been established in mammalian cells, such as probes for ATP^45^, acetylcholine^46^, glycine^24^, and NAD^+^/NADH^47^. Several genetically encoded fluorescent metabolite sensors, like the NAD^+^/NADH probe SoNar, have been successfully applied in the screening of anti-tumor agents^47^. l-2-HG has been exploited as a potential therapeutic target in renal cancer^14^ or a biomarker for cancer diagnosis and prognostic assessment^15,48^. The l-2-HG biosensors might also be utilized in the diagnosis and screening of anti-tumor agents for l-2-HG-related cancer.

In summary, a regulatory protein LhgR, which is involved in l-2-HG catabolism and specifically responds to l-2-HG, was identified in *P. putida* W619. Then, two FRET-based l-2-HG biosensors, LHGFR_0N3C_ and LHGFR_0N7C_, with excellent sensitivity, specificity, and stability, were constructed. The methods for quantitative estimation of l-2-HG concentrations in various biological samples and living cells by using l-2-HG biosensors were also established. We expect these l-2-HG biosensors to be of practical interest in future research on metabolism of l-2-HG and the diagnosis and treatment of l-2-HG-related diseases.

## Methods

Methods, including statements of data availability and any associated accession codes and references, are available in the online version of the paper.

## Acknowledgements

This work was supported by the grants of National Key R&D Program of China (2019YFA0904800, 2018YFA0901200, and 2019YFA0904900), the National Natural Science Foundation of China (31670041, 31970055), Shandong Provincial Funds for Distinguished Young Scientists (JQ 201806), Key R&D Program of Shandong Provincial (2019GSF107034, 2019GSF107039) and Qilu Young Scholar of Shandong University. The funders had no role in study design, data collection and interpretation, or the decision to submit the work for publication. We also thank Dr. Zhifeng Li and Dr. Jingyao Qu from Core Facilities for Life and Environmental Sciences (State Key Laboratory of Microbial Technology, Shandong University) for assistance in mass spectrographic analysis.

## Author contributions

C.G., C.M., and P.X. designed the research. Z.K., M.Z., K.G., Y.L., D.X., W.Z., and S.G. performed the research. Z.K. and C.G. analyzed the data. Z.K., C.G., C.M., and P.X. wrote the paper.

## Additional information

Any supplementary information and source data are available in the online version of the paper. Reprints and permissions information is available online at http://www.nature.com/reprints/index.html. Correspondence and requests for materials should be addressed to C.G. and P.X..

## Online methods

### Bacterial strains and culture conditions

The bacterial strains used in this study are listed in Supplementary Table 3. *E. coli* and its derivatives were cultured in Luria–Bertani (LB) broth at 37 °C and 180 rpm. *P. putida* KT2440 and its derivatives were grown in minimal salt mediums (MSMs) containing different carbon sources at 30 °C and 200 rpm. Antibiotics were used at the following concentrations: tetracycline at 30 μg mL^-1^; kanamycin at 50 μg mL^-1^; ampicillin at 100 μg mL^-1^; spectinomycin, at 50 μg mL^-1^; and chloramphenicol at 40 μg mL^-1^.

### Cloning of F2-*lhgR*-F1-*lhgO* and *lhgO*

All the plasmids and primers used in this study are listed in Supplementary Table 3 and Table 4, respectively. The gene segment F2-*lhgR*-F1-*lhgO* of *P. putida* W619 was synthesized by Tongyong Biosystem Co., Ltd (China). The *lhgO* gene of *P. putida* W619 was amplified and cloned into pME6032 plasmid using the restriction sites of EcoRI and KpnI to construct pME6032-*lhgO*, and the *P*_*tac*_ promoter of pME6032 was replaced by the gene segment F2-*lhgR*-F1-*lhgO* using the restriction sites of SacI and BamHI to construct pME6032-F2-*lhgR*-F1-*lhgO*, then both recombinant plasmids was transferred into different derivatives of *P. putida* KT2440 by electroporation, respectively.

### Construction of *P. putida* KT2440 and *E. coli* MG1655(DE3) mutants

Genes of *P. putida* KT2440 were deleted via allele exchange using the pK18mobsacB system^49^. Briefly, the homologous arms upstream and downstream of the target gene were PCR amplified and fused together by recombinant PCR. The generated fusion fragment was cloned into the suicide plasmid pK18*mobsacB*. The resulting plasmid was transferred into *P. putida* KT2440 by electroporation. The single crossover cells and the second crossover cells were sequentially screened from LB plates containing 50 μg mL^-1^ kanamycin or 10% (wt/vol) sucrose, respectively.

To construct the *E. coli* MG1655(DE3) (Δ*csiD*) mutant strain, the homologous arm upstream of the *csiD* gene, kanamycin resistance cassette, and the homologous arm downstream of the *csiD* gene were PCR amplified using the primers *csiD*-F1/*csiD*-R1, *csiD*-F2/*csiD*-R2, and *csiD*-F3/*csiD*-R3, respectively. The PCR products were fused together by recombinant PCR, and the resulting fusion was transferred into *E. coli* MG1655(DE3) harboring pTKRed plasmid following isopropyl-β-D-1-thiogalactopyranoside (IPTG) induction. The recombinant cells were selected on LB plates containing 50 μg mL^-1^ kanamycin at 37 °C. The pCP20 plasmid was transferred into the selected cells, followed by a second screening on LB plates containing 40 μg mL^-1^ chloramphenicol at 30 °C, then cultured in LB medium at 42 °C to eliminate pCP20 plasmid. The *lhgO* mutant of *E. coli* MG1655(DE3) were generated by the same process. All mutants were verified by PCR and sequencing.

### Enzymatic assay of LhgO

The derivatives of *P. putida* KT2440 were cultured in 50 mL MSMs with 5 g L^-1^ different compounds as carbon sources at 30 °C and 200 rpm. The cells were harvested at mid -log phase, washed twice and resuspended in phosphate-buffered saline (PBS), then lysed by sonication on ice after the addition of 1 mM phenylmethylsulfonyl fluoride (PMSF). The supernatants obtained were used for further enzyme activities measurements after a centrifugation process (13,000 × *g* for 10 min at 4 °C). Protein concentrations of the supernatants were determined using the Bradford protein assay kit (Sangon, China).

The activity of LhgO was assayed at 30 °C by monitored the reduction of dichlorophenol-indophenol (DCPIP) corresponding to the change of absorbance at 600 nm using a UV/visible spectrophotometer (Ultrospec 2100 pro, Amersham Biosciences, USA). The 800 μL reaction solution contained 0.1 mM l-2-HG, 0.05 mM DCPIP, 0.2 mM phenazine methosulfate (PMS) in PBS and 40 μL crude extracts. One unit of LhgO activity was defined as the amount of enzyme that catalyzed the reduction of 1 μmol of DCPIP per minute.

### Expression, purification, and characterization of LhgR

To express and purify the recombinant LhgR, the *lhgR* gene was PCR amplified using the primer pair *lhgR*-F/*lhgR*-R, which contained BamHI and HindIII restriction sites, respectively, and then cloned into the pETDuet-1 plasmid to construct pETDuet-*lhgR*. The *E. coli* BL21(DE3) strains harboring pETDuet-*lhgR* plasmid were grown to an OD_600_ of 0.6 in LB medium at 37 °C, after which the cells were induced for 12 h with 1 mM IPTG at 16 °C. The cells were harvested, washed twice and resuspended in buffer A (20 mM sodium phosphate and 500 mM sodium chloride, pH 7.4), then lysed by sonication on ice after addition of 1 mM PMSF and 10% (vol/vol) glycerol. The cell lysate was centrifuged at 13,000 × g for 40 min at 4 °C, and the resultant supernatant was loaded onto a HisTrap HP column (5 mL) equilibrated with buffer A. The target protein was eluted with buffer B (20 mM sodium phosphate, 500 mM sodium chloride, and 500 mM imidazole, pH 7.4), analyzed by 12.5% sodium dodecyl sulfate-polyacrylamide gel electrophoresis (SDS-PAGE), quantified by the Bradford protein assay kit (Sangon, China).

To determine the native molecular weight of LhgR, gel-filtration chromatography was performed using a Superdex 200 10/300 GL column (GE Healthcare, Germany) and standard proteins including thyroglobulin (669 kDa), ferritin (440 kDa), aldolase (158 kDa), conalbumin (75 kDa), ovalbumin (43 kDa), and ribonuclease A (13.7 kDa). The eluent buffer contained 50 mM sodium phosphate and 150 mM sodium chloride (pH 7.2).

### Electrophoretic mobility shift assays

Electrophoretic mobility shift assays (EMSAs) were carried out using the DNA fragment (F1 or F2) and purified LhgR. The DNA fragments were first amplified by primer pairs F1-F/F1-R and F2-F/F2-R, respectively. Then, either fragment at a concentration of 10 nM DNA was incubated with LhgR (0-160 nM) in 20 μL EMSA binding buffer (10 mM Tris-HCl [pH 7.4], 50 mM KCl, 0.5 mM EDTA, 10% [vol/vol] glycerol, and 1mM dithiothreitol [DTT]). The binding reactions were carried out at 30 °C for 30 min. Electrophoresis was performed on 6% native polyacrylamide gels at 4 °C and 170 V (constant voltage) for about 45 min, followed by staining with SYBR green I (TaKaRa, China) and photographing.

To characterize the effector of LhgR, purified LhgR was first incubated with L-lysine, 5-aminovalerate, glutarate, l-2-HG, D-2-HG, 2-KG, or succinate at 30 °C for 15 min, followed by incubation with the added DNA fragments at 30 °C for 30 min. The mixtures were subsequently subjected to electrophoresis.

### DNase I footprinting

DNase I footprinting assays were performed using the 6-carboxyfluorescein (FAM) labeled probe and purified LhgR. The DNA fragment F1 were PCR amplified using the primer pair F1-F/F1-R. The PCR products were cloned into pEASY-Blunt plasmid using pEASY-Blunt Cloning Kit (TransGen, China). The FAM-labeled probes were PCR amplified using the resulting plasmid and the primer pair M13F-FAM/M13R. Then, 350 ng probes were incubated with 2 μg purified LhgR in a total volume of 40 μL for 30 min at 30 °C. The DNase I digestion reaction was carried out by adding a total volume of 10 μL solution containing approximately 0.015 units of DNase I (Promega, USA) and 100 nmol CaCl_2_ and further incubating for 1 min at 37 °C, then stopped by adding a total volume of 140 μL stop solution containing 0.15% (wt/vol) SDS, 200 mM unbuffered sodium acetate, and 30 mM EDTA. The digested DNA fragments were first extracted with phenol-chloroform, then precipitated with ethanol and resuspended in 30 μL MiliQ water. The binding region of LhgR to DNA fragment F2 was analyzed using the same procedure.

### Construction and purification of LHGFR

The genes encoding mTFP and Venus were synthesized by Tongyong Biosystem Co., Ltd (China). The mTFP gene and Venus gene were amplified and cloned into pETDuet-1 plasmid using the BamHI and SacI restriction sites, and SalI and NotI restriction sites, respectively. Then either the full-length *lhgR* gene, its truncated variants, or variants with artificial linkers were inserted between mTFP and Venus by the T5 exonuclease DNA assembly (TEDA) method^50^, respectively. For expression in HEK293FT cells, the codon-optimized LHGFR_0N3C_ or LHGFR_0N7C_ sequence was synthesized and cloned into pcDNA3.1^(+)^ plasmid behind a Kozak sequence, 5’-GCCACC-3’. The l-2-HG biosensor LHGFR and its derivatives were expressed and purified using the same procedure.

### Characterization of LHGFR *in vitro*

Purified l-2-HG biosensors and different compounds were diluted by 50 mM Tris-HCl buffer (pH 7.4), mixed together in a black 96-well plate at a volume ratio of 3:1, and the fluorescence intensities were measured using an EnSight microplate reader (PerkinElmer, USA) with excitation at 430 nm, emission at 485 nm (mTFP) and 528 nm (Venus). The dose-response curves were fitted by OriginPro 2016 software (OriginLab) according to the following formula:

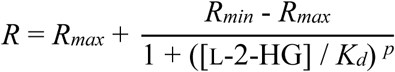

where *R, R*_*max*_, and *R*_*min*_ refer to the emission ratio of Venus to mTFP, ratio in the absence of l-2-HG, and ratio at saturation with l-2-HG, respectively. The [l-2-HG], *K*_*d*_, and *p* refer to the l-2-HG concentration, apparent dissociation constant, and Hill slope, respectively. Emission spectra were recorded at 430 nm excitation, in steps of 2 nm. The reversibility of LHGFR was determined by recording the emission ratios 10 min after the addition of 5 μM purified LhgO, the control test without addition of l-2-HG or purified LhgO was run in parallel. The pH stability of LHGFR was determined using 50 mM Tris-HCl buffer with pH adjusted from 4.0 to 9.0. The background fluorescence without the addition of LHGFR was subtracted.

In order to test the ability of LHGFR for quantitative analysis l-2-HG in different biological samples, purified LHGFR was diluted by 50 mM Tris-HCl (pH 7.4), while varying concentrations of l-2-HG was added into the serum and urine of healthy adults and bacteria culture medium and filtered through a 0.22 μm filter, respectively. The mixtures were then incubated in a black 96-well plate at a volume ratio of 3:1, and the emission ratios were determined. The background fluorescence without the addition of LHGFR was subtracted.

### Quantification of l-2-HG by HPLC and LC-MS/MS

When l-2-HG was used as carbon source to cultivate *P. putida* KT2440, its consumption was analyzed by using high-performance liquid chromatography (HPLC) system (Agilent 1100 series, Agilent Technologies, USA) equipped with an Aminex HPX-87H column (300 × 7.8 mm, Bio-Rad, USA) and a RID detector at 55 °C as described previously^7^.

To detect l-2-HG concentrations in various biological samples by liquid chromatography-tandem mass spectrometry (LC-MS/MS) system, the samples containing D,l-2-hydroxyglutarate disodium salt (2,3,3-D3) as internal standard (ITSD) were centrifuged at 13,000 × *g* for 15 min, then filtered through a 0.22 μm filter. The serum samples were mixed with methanol at a volume ratio of 1:3 and vortexed for 2 min to remove protein before centrifugation. Samples were analyzed by using a Thermo ultimate 3000 rapid separation liquid chromatograph system (ThermoFisher, USA) coupled with a Bruker impact HD ESI-Q-TOF mass spectrometer (Bruker Daltonics, Germany) in negative ion mode and equipped with a Chirobiotic R column (250 × 4.6 mm, Supelco Analytical, USA). Mobile phase was prepared from (A) 0.1% triethylamine adjusted to pH 4.5 with acetic acid or (B) methanol. The quantification was conducted with an injection volume of 20 μL, a constant 5% gradient of (B) at a flow rate of 0.5 mL min^-1^, and a total analysis time of 15 min.

### Characterization of LHGFR in living bacteria

*E. coli* BL21(DE3) strains harboring either pETDuet-LHGFR_0N3C_ or pETDuet-LHGFR_0N7C_ were grown to an OD_600_ of 0.6 in LB medium at 37 °C, after which the cells were induced overnight in the presence of 1 mM IPTG at 16 °C. The cultures were collected by centrifugation at 6000 × g for 5 min, washed three times, and resuspended to an OD_600_ of 2.5 by carbon starvation medium (MSM containing no carbon source) or glucose medium (MSM containing 20 mM glucose).

To characterize the sensitivity and specificity of LHGFR_0N3C_ and LHGFR_0N7C_ expressed in *E. coli* BL21(DE3), 90 μL cell suspensions following 8 h carbon starvation were mixed with 10 μL increasing concentrations of l-2-HG or other compounds, and then added into a black 96-well plate (total 100 μL/per well), the fluorescence intensities were determined using an EnSight microplate reader (PerkinElmer, USA) and the following instrument settings: excitation at 430 nm, emission at 485 nm (mTFP) and 528 nm (Venus), time intervals of 5 min, temperature at 37 °C, and shake at 180 rpm. For carbon starvation experiments, cell suspensions in carbon starvation medium or glucose medium were added into a black 96-well plate (100 μL/per well), then the fluorescence intensities were monitored every ten minutes. In order to analyze functions of CsiD and LhgO in endogenous l-2-HG anabolism and catabolism during carbon starvation, pETDuet-LHGFR_0N3C_ or pETDuet-LHGFR_0N7C_ was transferred into *E. coli* MG1655(DE3) and its variants, and the assays were performed using the same procedure. The background fluorescence from wells containing *E. coli* cultures harboring pETDuet-1 plasmid was subtracted at each emission wavelength.

### Cell culture and live-cell imaging

HEK293FT cells were cultured in high-glucose Dulbecco’s modified eagle medium (DMEM) supplemented with 10% (vol/vol) fetal bovine serum (FBS), 100 μnits mL^-1^ penicillin, and 100 μg mL^-1^ streptomycin (all purchased from ThermoFisher, USA), and kept at 37 °C in humidified air containing 5% CO_2_. For hypoxia experiments, cells were kept in a compact O_2_ and CO_2_ subchamber controller (ProOx C21, BioSpherix, USA) at 2% O_2_, 5% CO_2_, and balanced with N_2_ for 24 h. For construction of LHGFR expressing cell, HEK293FT cells were plated in 24-well plates and transfected with pcDNA3.1^(+)^ plasmid encoding either LHGFR_0N3C_ or LHGFR_0N7C_.

Live-cell imaging was carried out using a Zeiss 800 confocal microscope 48 h following transfection. LHGFR_0N3C_ or LHGFR_0N7C_ expressed in HEK293FT cells was excited using a 405 nm laser, and the emission was divided into a 460-500 nm channel (mTFP) and a 500-550 channel (Venus).

### Characterization of LHGFR in HEK293FT cells

To characterize the sensitivity and specificity of LHGFR expressed in HEK293FT cells, cells were trypsinized 48 h following transfection and suspended in 1 × Hank’s balanced salt solution supplemented with 20 mM HEPES. Increasing concentrations of l-2-HG or other compounds including glutarate, D-2-HG, and glucose was mixed with the cell suspensions in a 96-well plate, respectively. Digitonin at a concentration of 10 μM was used to induce cell permeabilization and deplete intracellular l-2-HG for *in vivo* response curves construction. Then, the fluorescence intensities were determined by a SpectraMax i3 fluorescence plate reader (Molecular Devices, USA) with excitation at 430 nm and emission at 485 nm (mTFP) and 528 nm (Venus). Basal l-2-HG concentration in HEK293FT cells under physiological conditions was determined by substituting the emission ratios of non-permeabilized HEK293FT cells into the calibrated *in vivo* response curves.

For the detection of hypoxia-induced production of l-2-HG, LHGFR_0N3C_ was expressed in HEK293FT cells and cultured for 24 h at 2% oxygen or 21% oxygen in the absence or presence of 5 mM dimethyl-2-ketoglutarate (DMαKG). The preparation of cell suspensions and the measurement of emission ratios were performed using the same procedure. The background fluorescence was subtracted at each emission wavelength.

### siRNA experiments

The following Silencer Select siRNAs used in this study were purchased from ThermoFisher Scientific (USA): negative control (4390846), L2HGDH-a (s36692), L2HGDH-b (s36693), LDHA (s351), and MDH2 (s8622). To analyze L2HGDH functions in l-2-HG catabolism, siRNA targeting L2HGDH and pcDNA3.1^(+)^ plasmid encoding either LHGFR_0N3C_ or LHGFR_0N7C_ were mixed with Lipofectamine 3000 Transfection Reagent (ThermoFisher, USA) in Opti-MEM Reduced Serum Medium, and the lipoplexes prepared were transfected into HEK293FT cells according to the manufacturer’s protocol. The fluorescence intensities were measured by a SpectraMax i3 fluorescence plate reader 48 h following transfection. Similarly, HEK293FT cells were transfected by LHGFR_0N3C_ and siRNAs targeting LDHA and MDH2 separately or in combination. After transfection, cells were cultured sequentially under normoxic condition for 24 h and hypoxic condition for 24 h in the presence of 5 mM DMαKG, then the fluorescence intensities were measured. The cells cultured under normoxic condition in the presence of 5 mM DMαKG for 48 h were set as control. The background fluorescence was subtracted at each emission wavelength.

### Data availability

The data supporting the findings of this study are available within the article and its Supplementary Information files and from the corresponding authors on request.

