## Supplementary Information for "A fluorescent l-2-hydroxyglutarate biosensor"

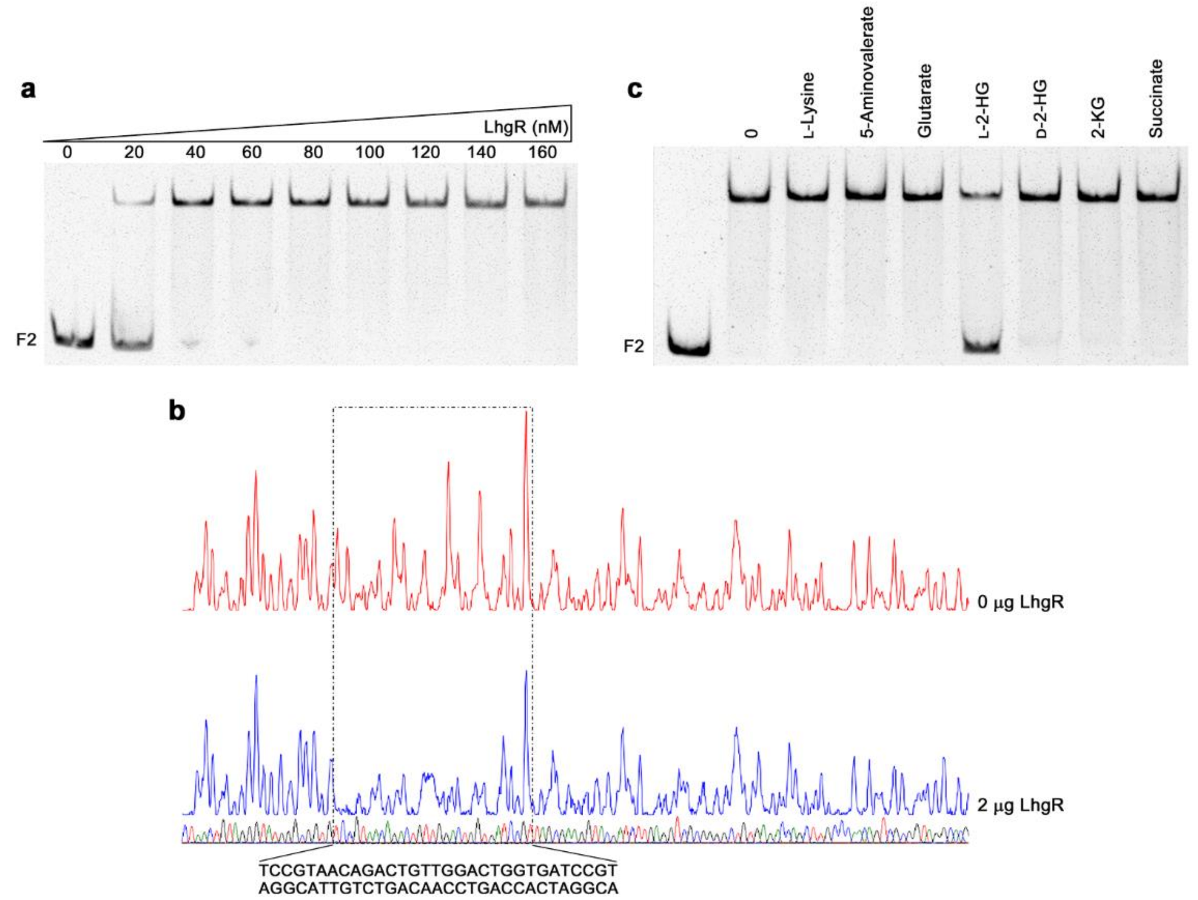


**Supplementary Figure 1** Analysis of the interaction between LhgR and the *lhgR* promoter region. **(a)** LhgR can bind to the *lhgR* promoter region. F2 fragment containing the *lhgR* promoter region (10 nM) was titrated by purified LhgR (0, 20, 40, 60, 80, 100, 120, 140, 160 nM). **(b)** DNase I footprinting analysis of LhgR binding to the *lhgR* promoter region. The F2 fragment was labeled with 6-carboxyfluorescein (FAM) and incubated with 2 µg LhgR (blue line) or without LhgR (red line). The region protected by LhgR is indicated with a dotted box. **(c)** L-2-HG prevent LhgR binding to the *lhgR* promoter region. EMSAs were carried out with F2 fragment (10 nM) and purified LhgR (70 nM) in the absence of any other tested compounds (0) and in the presence of 30 mM different compounds. The leftmost lane without LhgR was used as the control.

MVSKGEETTMGVIKPDMKIKLKMEGNVNGHAFVIEGEGEGKPYDGTNTINLE

VKEGAPLPFSYDILTTAFAYGNRAFTKYPDDIPNYFKQSFPEGYSWERTMTF

EDKGIVKVKSDISMEEDSFIYEIHLKGENFPPNGPVMQKKTTGWDASTERMY

VRDGVLKGDVKHKLLLEGGGHHRVDFKTIYRAKKAVKLPDYHFVDHRIEILN

HDKDYNKVTVYESAVARNSTDGMDELYKELMLELQRPDTLVERVVSAIRAEI

DSGRLAAEARLPTEQQLAEQLNVSRSVVREAVAQLKADGVLIARRGLGSYIS

KTPGGTVFRFPGSTGRKPDLVQMFEMRLWIETQAAAIAARRRDEHDLANMA

QALQEMLDKRSDFATASAADVAFHRAIAEASKNDYFVAFHDFLGGQLANAR

RTAWENSAAHSVGGSAEANREHQALYQAIADGDRQRAAACAEAHLRASAK

RLKIELPALDVDMVSKGEELFTGVVPILVELDGDVNGHKFSVSGEGEGDAT

YGKLTLKLICTTGKLPVPWPTLVTTLGYGLQCFARYPDHMKQHDFFKSAMP

EGYVQERTIFFKDDGNYKTRAEVKFEGDTLVNRIELKGIDFKEDGNILGHKLE

YNYNSHNVYITADKQKNGIKANFKIRHNIEDGGVQLADHYQQNTPIGDGPVL

LPDNHYLSYQSALSKDPNEKRDHMVLLEFVTAAGITLGMDELYK*

**Supplementary Figure 2** Full protein sequence of $LHGFR_{0N0C}$. The sequence of mTFP, LhgR, and Venus are highlighted by cyan, gray, and yellow, respectively. Truncation sites are indicated with dotted boxes. EL and VD are the amino acid sequences of the restriction sites SacI and SalI, respectively, and indicated with underlines.

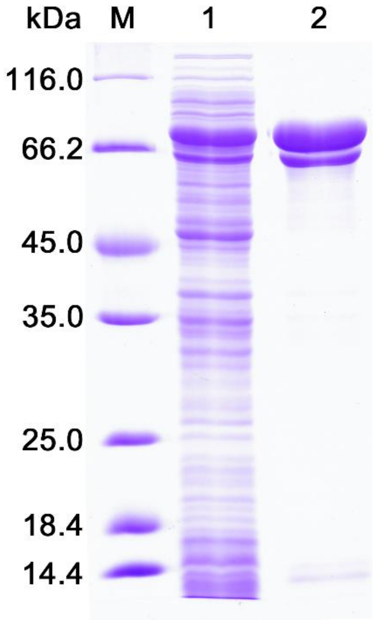


**Supplementary Figure 3** SDS-PAGE analysis of the purification of LHGFR$_{0N0C}$. Lane M, molecular weight markers; lane 1, crude extract of *E. coli* BL21(DE3) harboring pETDuet-LHGFR$_{0N0C}$; lane 2, purified His$_6$-tagged LHGFR$_{0N0C}$ using a HisTrap column.

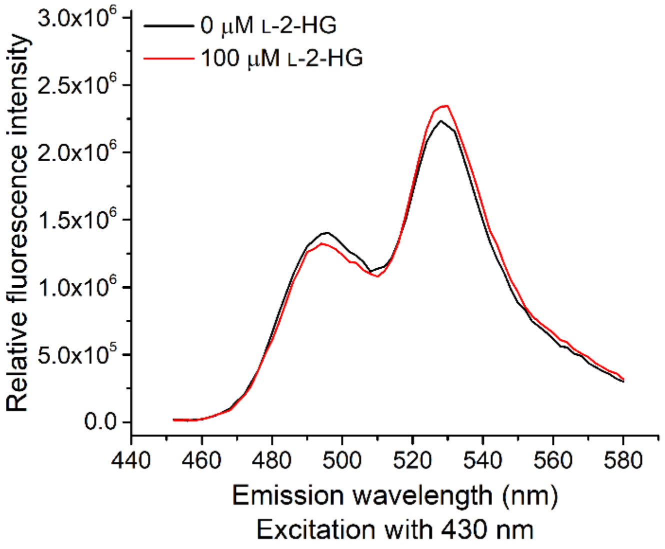


**Supplementary Figure 4** *In vitro* emission spectra analysis of LHGFR$_{0N0C}$. Fluorescence emission spectra changes of 1 μM LHGFR$_{0N0C}$ at 430 nm excitation with (red) or without (black) the addition of 100 μM L-2-HG were indicated.

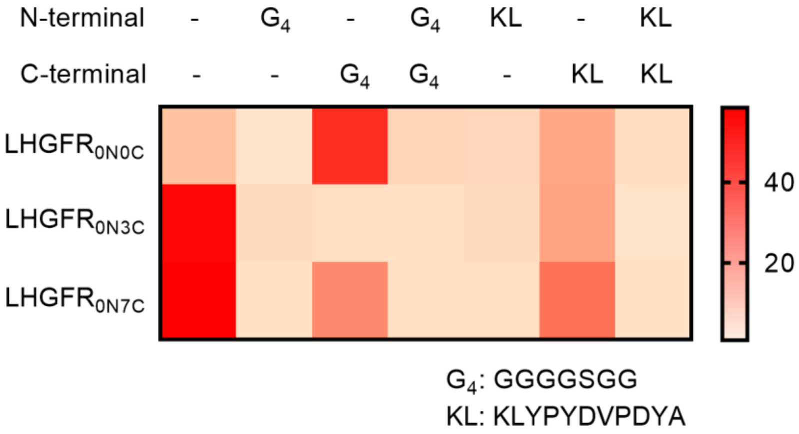


**Supplementary Figure 5** Heap map of $\Delta R_{max}$ to the LHGFR variants in which a series of artificial linkers were added between LhgR and fluorescent proteins. $G_4$ and KL indicate the flexible linker Gly-Gly-Gly-Gly-Ser-Gly-Gly and rigid linker Lys-Leu-Tyr-Pro-Tyr-Asp-Val-Pro-Asp-Tyr-Ala, respectively. Color indicates the value of $\Delta R_{max}$.

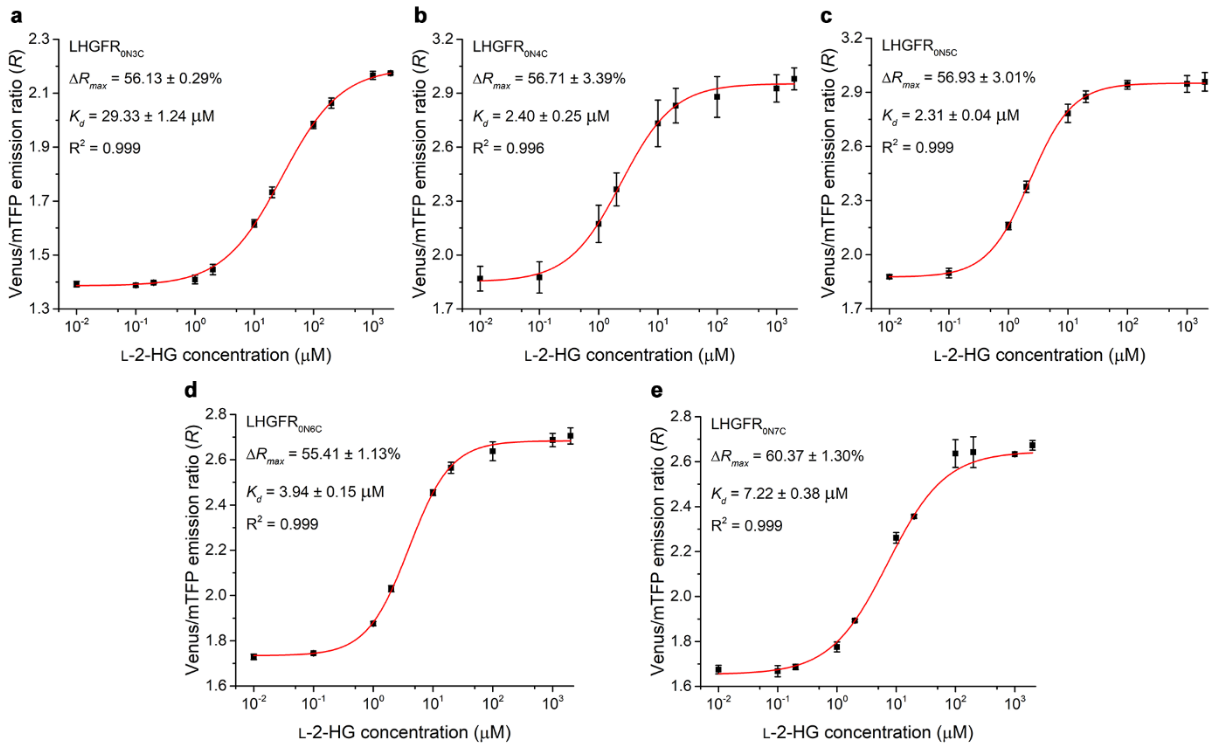


**Supplementary Figure 6** Dose-response curves of purified LHGFR variants truncated three to seven C-terminal amino acids in LhgR for L-2-HG. Dose-response curves of LHGFR$_{0N3C}$ **(a)**, LHGFR$_{0N4C}$ **(b)**, LHGFR$_{0N5C}$ **(c)**, LHGFR$_{0N6C}$ **(d)**, and LHGFR$_{0N7C}$ **(e)** for increasing concentrations (10 nM to 2 mM) of L-2-HG were indicated. The maximum ratio change ($\Delta R_{max}$), apparent dissociation constant ($K_d$), and $R^2$ were shown in each figure. All data shown are means ± s.d. (n = 3 independent experiments).

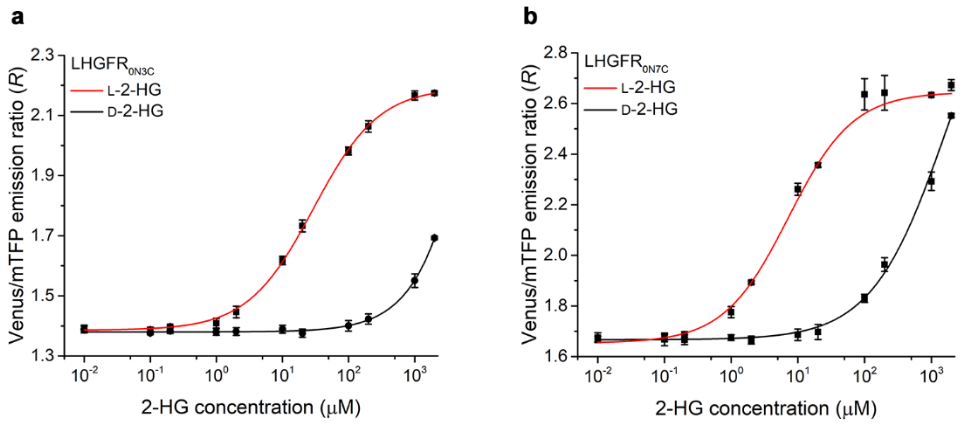


**Supplementary Figure 7** Comparison of the binding of L-2-HG and its mirror-image enantiomers D-2-HG with LHGFR$_{0N3C}$ **(a)** and LHGFR$_{0N7C}$ **(b)**. Dose-response curves of purified LHGFR$_{0N3C}$ **(a)** and LHGFR$_{0N7C}$ **(b)** for increasing concentrations (10 nM to 2 mM) of L-2-HG and D-2-HG were indicated. The dose-response curve for D-2-HG could not be fitted but the apparent $K_d$ of purified LHGFR$_{0N3C}$ in response to D-2-HG application was obviously on the order of millimoles. Purified LHGFR$_{0N7C}$ displayed a higher apparent $K_d$ in response to D-2-HG (1.85 ± 0.91 mM; black line) than L-2-HG (7.22 ± 0.38 μM; red line). All data shown are means ± s.d. (n = 3 independent experiments).

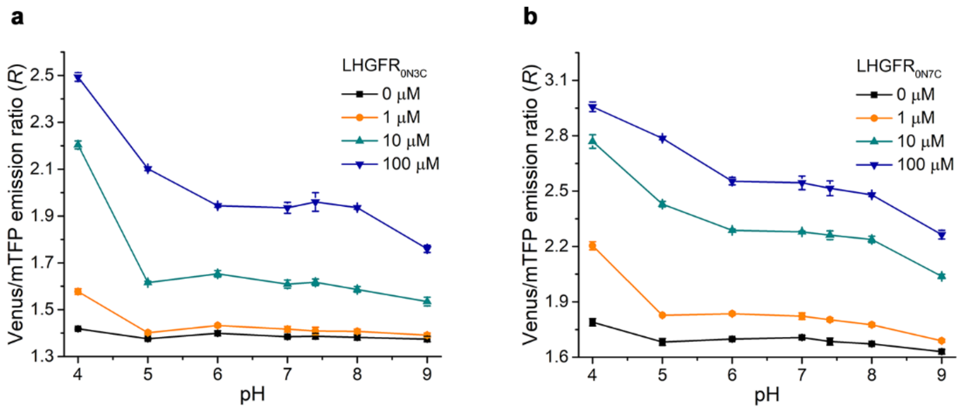


**Supplementary Figure 8** *In vitro* pH-stability analysis of purified LHGFR0N3C **(a)** and LHGFR0N7C **(b)**. Emission ratios of both biosensors in the presence of L-2-HG (0, 1, 10, and 100 μM) were determined at the indicated pH values. All data shown are means ± s.d. (n = 3 independent experiments).

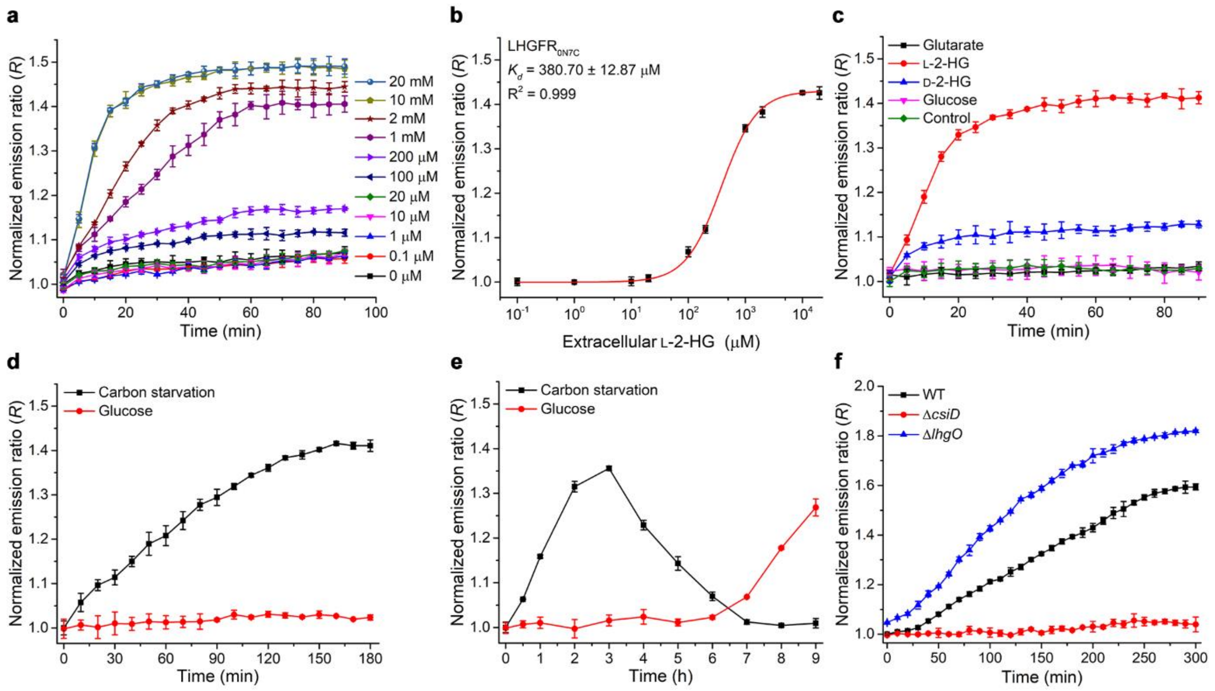


**Supplementary Figure 9** Monitoring L-2-HG fluctuations in living bacteria by LHGFR$_{0N7C}$. **(a)** Time course of the emission ratio changes of LHGFR$_{0N7C}$ expressed in *E. coli* BL21(DE3) in response to exogenous L-2-HG addition. All ratios were normalized to the control (ratio in the absence of L-2-HG at time point zero). **(b)** Normalized dose-response curve of LHGFR$_{0N7C}$ expressed in *E. coli* BL21(DE3) for increasing concentrations (100 nM to 20 mM) of L-2-HG at time point 60 min. **(c)** Time course of the emission ratio changes of LHGFR$_{0N7C}$ expressed in *E. coli* BL21(DE3) in response to the addition of 1 mM glutarate, L-2-HG, D-2-HG, or glucose. All data were normalized to the control (ratio in the absence of any tested compounds at time point zero). **(d)** Detection of carbon starvation-induced L-2-HG accumulation over time by LHGFR$_{0N7C}$ expressed in *E. coli* BL21(DE3). Emission ratio changes of LHGFR$_{0N7C}$ were measured when cultured in carbon starvation medium (black line) and medium with 20 mM glucose (red line). All data were normalized to samples under carbon starvation condition at time point zero. **(e)** Long-term detection of L-2-HG fluctuations by LHGFR$_{0N7C}$ expressed in *E. coli* BL21(DE3). All data were normalized to samples under carbon starvation condition at time point zero. **(f)** Identification of the roles of CsiD and LhgO in endogenous L-2-HG catabolism during carbon starvation by LHGFR$_{0N7C}$. Emission ratio changes of LHGFR$_{0N7C}$ expressed in *E. coli* MG1655(DE3) wild-type (black line), *E. coli* MG1655(DE3) (Δ*csiD*) (red line), and *E. coli* MG1655(DE3) (Δ*lhgO*) (blue line) were measured in carbon starvation medium. All data were normalized to time point zero of wild-type strain and shown as means ± s.d. (n = 3 independent experiments).

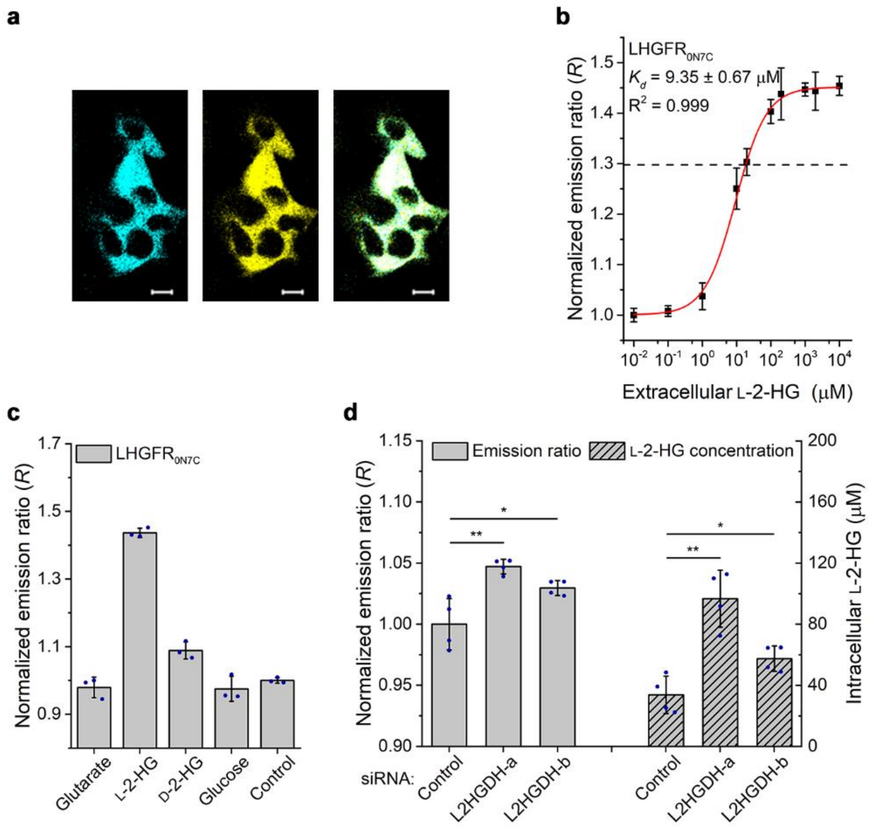


**Supplementary Figure 10** Monitoring L-2-HG fluctuations in human cells by LHGFR0N7C. **(a)** Confocal microscopy images of LHGFR0N7C-expressing HEK293FT cells. The images are represented as mTFP channel, Venus channel, and overlay channel from left to right. Scale bar, 10 µm. **(b)** Normalized dose-response curve of LHGFR0N7C expressed in HEK293FT cells with increasing concentrations (10 nM to 10 mM) of L-2-HG. Cells were permeabilized with 10 µM digitonin. The emission ratio of non-permeabilized HEK293FT cells under physiological conditions is indicated with black dash line. **(c)** Responses of LHGFR0N7C expressed in HEK293FT cells to exogenously added 1 mM glutarate, L-2-HG, D-2-HG, and glucose. All data were normalized to the control (ratio in the absence of any tested compounds). **(d)** Identification of the function of L2HGDH in L-2-HG catabolism by LHGFR0N7C. The emission ratio was measured after co-transfecting siRNA targeting L2HGDH and LHGFR0N7C for 48 h. Emission ratio was normalized to the control condition. All data shown are means ± s.d. (n = 3, 3, and 4 independent experiments for **b**, **c**, and **d**). *, $P < 0.05$ in two-tailed *t* test; **, $P < 0.01$ in two-tailed *t* test.

**Supplementary Table 1 Evaluation of the performance of LHGFR for quantification of L-2-HG in various biological samples.**

| Condition | Approach | Concentration (μM) | | | Accuracy (%)[a] | | | Precision (RSD%)[b] |
|---|---|---|---|---|---|---|---|---|
| | | Sample 1 | Sample 2 | Sample 3 | Sample 1 | Sample 2 | Sample 3 | |
| | Standard | 8 | 40 | 80 | 100 | 100 | 100 | |
| Serum | LC-MS/MS | 7.77 | 45.88 | 94.85 | 97.19 | 114.70 | 118.56 | 10.34 |
| | LHGFR$_{0N3C}$ | 9.11 | 42.75 | 94.40 | 113.88 | 106.88 | 117.99 | 4.98 |
| | LHGFR$_{0N7C}$ | 8.92 | 38.19 | 91.50 | 111.55 | 95.48 | 114.38 | 9.51 |
| Urine | LC-MS/MS | 8.53 | 38.65 | 84.62 | 106.63 | 96.62 | 105.77 | 5.39 |
| | LHGFR$_{0N3C}$ | 9.49 | 46.81 | 89.36 | 118.62 | 117.02 | 111.71 | 3.12 |
| | LHGFR$_{0N7C}$ | 8.45 | 45.77 | 92.79 | 105.63 | 114.42 | 115.99 | 4.98 |
| Bacterial culture medium | LC-MS/MS | 8.02 | 50.08 | 88.33 | 100.31 | 125.20 | 110.41 | 11.18 |
| | LHGFR$_{0N3C}$ | 8.04 | 38.73 | 79.83 | 100.46 | 96.81 | 99.79 | 1.96 |
| | LHGFR$_{0N7C}$ | 8.36 | 42.28 | 80.71 | 104.47 | 105.69 | 100.88 | 2.41 |

[a]Accuracy% = $\frac{\text{Concentration determined by LC-MS/MS or LHGFR}}{\text{Defined concentration}}$.

[b]Precision% = $\frac{\text{Standard derivation of accuracy}}{\text{Mean value of accuracy}}$.

**Supplementary Table 2 LOD and detection range of various methods for quantification of L-2-HG in various biological samples.**

| Approach | Serum | | Urine | | Bacterial culture medium | |
|---|---|---|---|---|---|---|
| | LOD (μM)[a] | Detection range (μM) | LOD (μM)[a] | Detection range (μM) | LOD (μM)[a] | Detection range (μM) |
| HPLC | 400 | > 400 | 100 | > 100 | 100 | > 100 |
| LC-MS/MS | 4 | > 4 | 1 | > 1 | 1 | > 1 |
| $LHGFR_{0N3C}$ | 5.84 | 5.84 - 4000 | 15.74 | 15.74 - 4000 | 5.48 | 5.48 - 4000 |
| $LHGFR_{0N7C}$ | 1.68 | 1.68 - 400 | 0.92 | 0.92 - 400 | 0.82 | 0.82 - 400 |

[a]The limits of detection (LOD) is calculated by interpolating the average background counts + 3 × standard deviation value.

**Supplementary Table 3 Strains and plasmids used in this study**

| Strain or plasmid[a] | Relevant characteristics[b] |
|---|---|
| **Strain** | |
| *P. putida* KT2440 | Wild-type |
| *P. putida* KT2440 (Δ*lhgO*) | *P. putida* KT2440 with a deletion of the *lhgO* gene |
| *P. putida* KT2440 (Δ*csiR*Δ*lhgO*) | *P. putida* KT2440 with deletions of the *csiR* gene and *lhgO* gene |
| *P. putida* KT2440 (Δ*csiR*Δ*csiD*Δ*lhgO*) | *P. putida* KT2440 with deletions of the *csiR* gene, *csiD* gene, and *lhgO* gene |
| *P. putida* KT2440 (Δ*lhgO*)-pME6032 | *P. putida* KT2440 (Δ*lhgO*) harboring the empty plasmid pME6032 |
| *P. putida* KT2440 (Δ*lhgO*)-pME6032-*lhgO* | *P. putida* KT2440 (Δ*lhgO*) harboring the plasmid pME6032-*lhgO* |
| *P. putida* KT2440 (Δ*csiR*Δ*lhgO*)-pME6032 | *P. putida* KT2440 (Δ*csiR*Δ*lhgO*) harboring the empty plasmid pME6032 |
| *P. putida* KT2440 (Δ*csiR*Δ*lhgO*)-pME6032-F2-*lhgR*-F1-*lhgO* | *P. putida* KT2440 (Δ*csiR*Δ*lhgO*) harboring the plasmid pME6032-F2-*lhgR*-F1-*lhgO* |
| *P. putida* KT2440 (Δ*csiR*Δ*csiD*Δ*lhgO*)-pME6032 | *P. putida* KT2440 (Δ*csiR*Δ*csiD*Δ*lhgO*) harboring the empty plasmid pME6032 |
| *P. putida* KT2440 (Δ*csiR*Δ*csiD*Δ*lhgO*)-pME6032-F2-*lhgR*-F1-*lhgO* | *P. putida* KT2440 (Δ*csiR*Δ*csiD*Δ*lhgO*) harboring the plasmid pME6032-F2-*lhgR*-F1-*lhgO* |
| *E. coli* DH5α | $F^{-}$ φ80*lacZ*ΔM15 Δ(*lacZYA*-*argF*)U169 *deoR recA*1 *endA*1 *hsdR*17($r_K^{-}$, $m_K^{+}$) *phoA supE*44 $\lambda^{-}$ *thi*-1 *gyrA*96 *relA*1 |
| *E. coli* BL21(DE3) | $F^{-}$ *ompT hsdSB*(*rB*- *mB*-) *gal*(λ *c I* 857 *ind1 Sam*7 *nin*5 *lac*UV5-T7*gene*1) *dcm* (DE3) |
| *E. coli* BL21(DE3)-LhgR | *E. coli* BL21(DE3) harboring the expression plasmid pETDuet-*lhgR* |
| *E. coli* BL21(DE3)-$LHGFR_{0N0C}$ | *E. coli* BL21(DE3) harboring the expression plasmid pETDuet-$LHGFR_{0N0C}$ |
| *E. coli* BL21(DE3)-$LHGFR_{0N1C}$ | *E. coli* BL21(DE3) harboring the expression plasmid pETDuet-$LHGFR_{0N1C}$ |
| *E. coli* BL21(DE3)-$LHGFR_{0N2C}$ | *E. coli* BL21(DE3) harboring the expression plasmid pETDuet-$LHGFR_{0N2C}$ |

| | |
|---|---|
| *E. coli* BL21(DE3)-LHGFR$_{0N3C}$ | *E. coli* BL21(DE3) harboring the expression plasmid pETDuet-LHGFR$_{0N3C}$ |
| *E. coli* BL21(DE3)-LHGFR$_{0N4C}$ | *E. coli* BL21(DE3) harboring the expression plasmid pETDuet-LHGFR$_{0N4C}$ |
| *E. coli* BL21(DE3)-LHGFR$_{0N5C}$ | *E. coli* BL21(DE3) harboring the expression plasmid pETDuet-LHGFR$_{0N5C}$ |
| *E. coli* BL21(DE3)-LHGFR$_{0N6C}$ | *E. coli* BL21(DE3) harboring the expression plasmid pETDuet-LHGFR$_{0N6C}$ |
| *E. coli* BL21(DE3)-LHGFR$_{0N7C}$ | *E. coli* BL21(DE3) harboring the expression plasmid pETDuet-LHGFR$_{0N7C}$ |
| *E. coli* BL21(DE3)-LHGFR$_{0N8C}$ | *E. coli* BL21(DE3) harboring the expression plasmid pETDuet-LHGFR$_{0N8C}$ |
| *E. coli* BL21(DE3)-LHGFR$_{0N9C}$ | *E. coli* BL21(DE3) harboring the expression plasmid pETDuet-LHGFR$_{0N9C}$ |
| *E. coli* BL21(DE3)-LHGFR$_{0N10C}$ | *E. coli* BL21(DE3) harboring the expression plasmid pETDuet-LHGFR$_{0N10C}$ |
| *E. coli* BL21(DE3)-LHGFR$_{0N11C}$ | *E. coli* BL21(DE3) harboring the expression plasmid pETDuet-LHGFR$_{0N11C}$ |
| *E. coli* BL21(DE3)-LHGFR$_{0N12C}$ | *E. coli* BL21(DE3) harboring the expression plasmid pETDuet-LHGFR$_{0N12C}$ |
| *E. coli* BL21(DE3)-LHGFR$_{0N13C}$ | *E. coli* BL21(DE3) harboring the expression plasmid pETDuet-LHGFR$_{0N13C}$ |
| *E. coli* BL21(DE3)-LHGFR$_{0N14C}$ | *E. coli* BL21(DE3) harboring the expression plasmid pETDuet-LHGFR$_{0N14C}$ |
| *E. coli* BL21(DE3)-LHGFR$_{0N15C}$ | *E. coli* BL21(DE3) harboring the expression plasmid pETDuet-LHGFR$_{0N15C}$ |
| *E. coli* BL21(DE3)-LHGFR$_{1N0C}$ | *E. coli* BL21(DE3) harboring the expression plasmid pETDuet-LHGFR$_{1N0C}$ |
| *E. coli* BL21(DE3)-LHGFR$_{1N1C}$ | *E. coli* BL21(DE3) harboring the expression plasmid pETDuet-LHGFR$_{1N1C}$ |
| *E. coli* BL21(DE3)-LHGFR$_{1N2C}$ | *E. coli* BL21(DE3) harboring the expression plasmid pETDuet-LHGFR$_{1N2C}$ |
| *E. coli* BL21(DE3)-LHGFR$_{1N3C}$ | *E. coli* BL21(DE3) harboring the expression plasmid pETDuet-LHGFR$_{1N3C}$ |
| *E. coli* BL21(DE3)-LHGFR$_{1N4C}$ | *E. coli* BL21(DE3) harboring the expression plasmid pETDuet-LHGFR$_{1N4C}$ |
| *E. coli* BL21(DE3)-LHGFR$_{1N5C}$ | *E. coli* BL21(DE3) harboring the expression plasmid pETDuet-LHGFR$_{1N5C}$ |
| *E. coli* BL21(DE3)-LHGFR$_{1N6C}$ | *E. coli* BL21(DE3) harboring the expression plasmid pETDuet-LHGFR$_{1N6C}$ |
| *E. coli* BL21(DE3)-LHGFR$_{1N7C}$ | *E. coli* BL21(DE3) harboring the expression plasmid pETDuet-LHGFR$_{1N7C}$ |
| *E. coli* BL21(DE3)-LHGFR$_{2N0C}$ | *E. coli* BL21(DE3) harboring the expression plasmid pETDuet-LHGFR$_{2N0C}$ |

| | |
|---|---|
| *E. coli* BL21(DE3)-LHGFR$_{2N1C}$ | *E. coli* BL21(DE3) harboring the expression plasmid pETDuet-LHGFR$_{2N1C}$ |
| *E. coli* BL21(DE3)-LHGFR$_{2N2C}$ | *E. coli* BL21(DE3) harboring the expression plasmid pETDuet-LHGFR$_{2N2C}$ |
| *E. coli* BL21(DE3)-LHGFR$_{2N3C}$ | *E. coli* BL21(DE3) harboring the expression plasmid pETDuet-LHGFR$_{2N3C}$ |
| *E. coli* BL21(DE3)-LHGFR$_{2N4C}$ | *E. coli* BL21(DE3) harboring the expression plasmid pETDuet-LHGFR$_{2N4C}$ |
| *E. coli* BL21(DE3)-LHGFR$_{2N5C}$ | *E. coli* BL21(DE3) harboring the expression plasmid pETDuet-LHGFR$_{2N5C}$ |
| *E. coli* BL21(DE3)-LHGFR$_{2N6C}$ | *E. coli* BL21(DE3) harboring the expression plasmid pETDuet-LHGFR$_{2N6C}$ |
| *E. coli* BL21(DE3)-LHGFR$_{2N7C}$ | *E. coli* BL21(DE3) harboring the expression plasmid pETDuet-LHGFR$_{2N7C}$ |
| *E. coli* BL21(DE3)-LHGFR$_{3N0C}$ | *E. coli* BL21(DE3) harboring the expression plasmid pETDuet-LHGFR$_{3N0C}$ |
| *E. coli* BL21(DE3)-LHGFR$_{3N1C}$ | *E. coli* BL21(DE3) harboring the expression plasmid pETDuet-LHGFR$_{3N1C}$ |
| *E. coli* BL21(DE3)-LHGFR$_{3N2C}$ | *E. coli* BL21(DE3) harboring the expression plasmid pETDuet-LHGFR$_{3N2C}$ |
| *E. coli* BL21(DE3)-LHGFR$_{3N3C}$ | *E. coli* BL21(DE3) harboring the expression plasmid pETDuet-LHGFR$_{3N3C}$ |
| *E. coli* BL21(DE3)-LHGFR$_{3N4C}$ | *E. coli* BL21(DE3) harboring the expression plasmid pETDuet-LHGFR$_{3N4C}$ |
| *E. coli* BL21(DE3)-LHGFR$_{3N5C}$ | *E. coli* BL21(DE3) harboring the expression plasmid pETDuet-LHGFR$_{3N5C}$ |
| *E. coli* BL21(DE3)-LHGFR$_{3N6C}$ | *E. coli* BL21(DE3) harboring the expression plasmid pETDuet-LHGFR$_{3N6C}$ |
| *E. coli* BL21(DE3)-LHGFR$_{3N7C}$ | *E. coli* BL21(DE3) harboring the expression plasmid pETDuet-LHGFR$_{3N7C}$ |
| *E. coli* BL21(DE3)-LHGFR$_{4N0C}$ | *E. coli* BL21(DE3) harboring the expression plasmid pETDuet-LHGFR$_{4N0C}$ |
| *E. coli* BL21(DE3)-LHGFR$_{4N1C}$ | *E. coli* BL21(DE3) harboring the expression plasmid pETDuet-LHGFR$_{4N1C}$ |
| *E. coli* BL21(DE3)-LHGFR$_{4N2C}$ | *E. coli* BL21(DE3) harboring the expression plasmid pETDuet-LHGFR$_{4N2C}$ |
| *E. coli* BL21(DE3)-LHGFR$_{4N3C}$ | *E. coli* BL21(DE3) harboring the expression plasmid pETDuet-LHGFR$_{4N3C}$ |
| *E. coli* BL21(DE3)-LHGFR$_{4N4C}$ | *E. coli* BL21(DE3) harboring the expression plasmid pETDuet-LHGFR$_{4N4C}$ |
| *E. coli* BL21(DE3)-LHGFR$_{4N5C}$ | *E. coli* BL21(DE3) harboring the expression plasmid pETDuet-LHGFR$_{4N5C}$ |
| *E. coli* BL21(DE3)-LHGFR$_{4N6C}$ | *E. coli* BL21(DE3) harboring the expression plasmid pETDuet-LHGFR$_{4N6C}$ |

| | |
|---|---|
| *E. coli* BL21(DE3)-LHGFR$_{4N7C}$ | *E. coli* BL21(DE3) harboring the expression plasmid pETDuet-LHGFR$_{4N7C}$ |
| *E. coli* BL21(DE3)-LHGFR$_{G4\text{-}0N0C}$ | *E. coli* BL21(DE3) harboring the expression plasmid pETDuet-LHGFR$_{G4\text{-}0N0C}$ |
| *E. coli* BL21(DE3)-LHGFR$_{0N0C\text{-}G4}$ | *E. coli* BL21(DE3) harboring the expression plasmid pETDuet-LHGFR$_{0N0C\text{-}G4}$ |
| *E. coli* BL21(DE3)-LHGFR$_{G4\text{-}0N0C\text{-}G4}$ | *E. coli* BL21(DE3) harboring the expression plasmid pETDuet-LHGFR$_{G4\text{-}0N0C\text{-}G4}$ |
| *E. coli* BL21(DE3)-LHGFR$_{KL\text{-}0N0C}$ | *E. coli* BL21(DE3) harboring the expression plasmid pETDuet-LHGFR$_{KL\text{-}0N0C}$ |
| *E. coli* BL21(DE3)-LHGFR$_{0N0C\text{-}KL}$ | *E. coli* BL21(DE3) harboring the expression plasmid pETDuet-LHGFR$_{0N0C\text{-}KL}$ |
| *E. coli* BL21(DE3)-LHGFR$_{KL\text{-}0N0C\text{-}KL}$ | *E. coli* BL21(DE3) harboring the expression plasmid pETDuet-LHGFR$_{KL\text{-}0N0C\text{-}KL}$ |
| *E. coli* BL21(DE3)-LHGFR$_{G4\text{-}0N3C}$ | *E. coli* BL21(DE3) harboring the expression plasmid pETDuet-LHGFR$_{G4\text{-}0N3C}$ |
| *E. coli* BL21(DE3)-LHGFR$_{0N3C\text{-}G4}$ | *E. coli* BL21(DE3) harboring the expression plasmid pETDuet-LHGFR$_{0N3C\text{-}G4}$ |
| *E. coli* BL21(DE3)-LHGFR$_{G4\text{-}0N3C\text{-}G4}$ | *E. coli* BL21(DE3) harboring the expression plasmid pETDuet-LHGFR$_{G4\text{-}0N3C\text{-}G4}$ |
| *E. coli* BL21(DE3)-LHGFR$_{KL\text{-}0N3C}$ | *E. coli* BL21(DE3) harboring the expression plasmid pETDuet-LHGFR$_{KL\text{-}0N3C}$ |
| *E. coli* BL21(DE3)-LHGFR$_{0N3C\text{-}KL}$ | *E. coli* BL21(DE3) harboring the expression plasmid pETDuet-LHGFR$_{0N3C\text{-}KL}$ |
| *E. coli* BL21(DE3)-LHGFR$_{KL\text{-}0N3C\text{-}KL}$ | *E. coli* BL21(DE3) harboring the expression plasmid pETDuet-LHGFR$_{KL\text{-}0N3C\text{-}KL}$ |
| *E. coli* BL21(DE3)-LHGFR$_{G4\text{-}0N7C}$ | *E. coli* BL21(DE3) harboring the expression plasmid pETDuet-LHGFR$_{G4\text{-}0N7C}$ |
| *E. coli* BL21(DE3)-LHGFR$_{0N7C\text{-}G4}$ | *E. coli* BL21(DE3) harboring the expression plasmid pETDuet-LHGFR$_{0N7C\text{-}G4}$ |
| *E. coli* BL21(DE3)-LHGFR$_{G4\text{-}0N7C\text{-}G4}$ | *E. coli* BL21(DE3) harboring the expression plasmid pETDuet-LHGFR$_{G4\text{-}0N7C\text{-}G4}$ |
| *E. coli* BL21(DE3)-LHGFR$_{KL\text{-}0N7C}$ | *E. coli* BL21(DE3) harboring the expression plasmid pETDuet-LHGFR$_{KL\text{-}0N7C}$ |
| *E. coli* BL21(DE3)-LHGFR$_{0N7C\text{-}KL}$ | *E. coli* BL21(DE3) harboring the expression plasmid pETDuet-LHGFR$_{0N7C\text{-}KL}$ |
| *E. coli* BL21(DE3)-LHGFR$_{KL\text{-}0N7C\text{-}KL}$ | *E. coli* BL21(DE3) harboring the expression plasmid pETDuet-LHGFR$_{KL\text{-}0N7C\text{-}KL}$ |
| *E. coli* MG1655 | Wild-type |
| *E. coli* MG1655(DE3) | *E. coli* MG1655 expressing T7 RNA polymerase |
| *E. coli* MG1655(DE3) (Δ*csiD*) | *E. coli* MG1655(DE3) with a deletion of the *csiD* gene |

| | |
|---|---|
| *E. coli* MG1655(DE3) (Δ*lhgO*) | *E. coli* MG1655(DE3) with a deletion of the *lhgO* gene |
| *E. coli* MG1655(DE3)-LHGFR$_{0N3C}$ | *E. coli* MG1655(DE3) harboring the expression plasmid pETDuet-LHGFR$_{0N3C}$ |
| *E. coli* MG1655(DE3)-LHGFR$_{0N7C}$ | *E. coli* MG1655(DE3) harboring the expression plasmid pETDuet-LHGFR$_{0N7C}$ |
| *E. coli* MG1655(DE3) (Δ*csiD*)-LHGFR$_{0N3C}$ | *E. coli* MG1655(DE3) (Δ*csiD*) harboring the expression plasmid pETDuet-LHGFR$_{0N3C}$ |
| *E. coli* MG1655(DE3) (Δ*csiD*)-LHGFR$_{0N7C}$ | *E. coli* MG1655(DE3) (Δ*csiD*) harboring the expression plasmid pETDuet-LHGFR$_{0N7C}$ |
| *E. coli* MG1655(DE3) (Δ*lhgO*)-LHGFR$_{0N3C}$ | *E. coli* MG1655(DE3) (Δ*lhgO*) harboring the expression plasmid pETDuet-LHGFR$_{0N3C}$ |
| *E. coli* MG1655(DE3) (Δ*lhgO*)-LHGFR$_{0N7C}$ | *E. coli* MG1655(DE3) (Δ*lhgO*) harboring the expression plasmid pETDuet-LHGFR$_{0N7C}$ |
| **Plasmid** | |
| pME6032 | pVS1-p15A *E. coli*-*Pseudomonas* shuttle vector, *lacI*$^q$-*P*$_{tac}$ expression vector; Tc$^r$ |
| pME6032-*lhgO* | pME6032 contained *lhgO* gene of *P. putida* W619 |
| pME6032-F2-*lhgR*-F1-*lhgO* | pME6032 contained F2-*lhgR*-F1-*lhgO* gene segment of *P. putida* W619 and the promoter *P*$_{tac}$ was deleted |
| pK18*mobsacB* | Suicide plasmid for gene knockout; Km$^r$ |
| pK18*mobsacB*-Δ*lhgO* | Partial lengths of *lhgO* were inserted into pK18*mobsacB* |
| pK18*mobsacB*-Δ*csiR* | Partial lengths of *csiR* were inserted into pK18*mobsacB* |
| pK18*mobsacB*-Δ*csiD* | Partial lengths of *csiD* were inserted into pK18*mobsacB* |
| pETDuet-1 | Vector for protein expression; Ap$^r$ |
| pETDuet-*lhgR* | pETDuet-1 contained *lhgR* gene of *P. putida* W619 |
| pETDuet-LHGFR$_{0N0C}$ | pETDuet-1 contained the gene of LHGFR$_{0N0C}$ |
| pETDuet-LHGFR$_{0N1C}$ | pETDuet-1 contained the gene of LHGFR$_{0N1C}$ |
| pETDuet-LHGFR$_{0N2C}$ | pETDuet-1 contained the gene of LHGFR$_{0N2C}$ |
| pETDuet-LHGFR$_{0N3C}$ | pETDuet-1 contained the gene of LHGFR$_{0N3C}$ |

| | |
|---|---|
| pETDuet-LHGFR$_{0N4C}$ | pETDuet-1 contained the gene of LHGFR$_{0N4C}$ |
| pETDuet-LHGFR$_{0N5C}$ | pETDuet-1 contained the gene of LHGFR$_{0N5C}$ |
| pETDuet-LHGFR$_{0N6C}$ | pETDuet-1 contained the gene of LHGFR$_{0N6C}$ |
| pETDuet-LHGFR$_{0N7C}$ | pETDuet-1 contained the gene of LHGFR$_{0N7C}$ |
| pETDuet-LHGFR$_{0N8C}$ | pETDuet-1 contained the gene of LHGFR$_{0N8C}$ |
| pETDuet-LHGFR$_{0N9C}$ | pETDuet-1 contained the gene of LHGFR$_{0N9C}$ |
| pETDuet-LHGFR$_{0N10C}$ | pETDuet-1 contained the gene of LHGFR$_{0N10C}$ |
| pETDuet-LHGFR$_{0N11C}$ | pETDuet-1 contained the gene of LHGFR$_{0N11C}$ |
| pETDuet-LHGFR$_{0N12C}$ | pETDuet-1 contained the gene of LHGFR$_{0N12C}$ |
| pETDuet-LHGFR$_{0N13C}$ | pETDuet-1 contained the gene of LHGFR$_{0N13C}$ |
| pETDuet-LHGFR$_{0N14C}$ | pETDuet-1 contained the gene of LHGFR$_{0N14C}$ |
| pETDuet-LHGFR$_{0N15C}$ | pETDuet-1 contained the gene of LHGFR$_{0N15C}$ |
| pETDuet-LHGFR$_{1N0C}$ | pETDuet-1 contained the gene of LHGFR$_{1N0C}$ |
| pETDuet-LHGFR$_{1N1C}$ | pETDuet-1 contained the gene of LHGFR$_{1N1C}$ |
| pETDuet-LHGFR$_{1N2C}$ | pETDuet-1 contained the gene of LHGFR$_{1N2C}$ |
| pETDuet-LHGFR$_{1N3C}$ | pETDuet-1 contained the gene of LHGFR$_{1N3C}$ |
| pETDuet-LHGFR$_{1N4C}$ | pETDuet-1 contained the gene of LHGFR$_{1N4C}$ |
| pETDuet-LHGFR$_{1N5C}$ | pETDuet-1 contained the gene of LHGFR$_{1N5C}$ |
| pETDuet-LHGFR$_{1N6C}$ | pETDuet-1 contained the gene of LHGFR$_{1N6C}$ |
| pETDuet-LHGFR$_{1N7C}$ | pETDuet-1 contained the gene of LHGFR$_{1N7C}$ |
| pETDuet-LHGFR$_{2N0C}$ | pETDuet-1 contained the gene of LHGFR$_{2N0C}$ |
| pETDuet-LHGFR$_{2N1C}$ | pETDuet-1 contained the gene of LHGFR$_{2N1C}$ |

| | |
|---|---|
| pETDuet-$LHGFR_{2N2C}$ | pETDuet-1 contained the gene of $LHGFR_{2N2C}$ |
| pETDuet-$LHGFR_{2N3C}$ | pETDuet-1 contained the gene of $LHGFR_{2N3C}$ |
| pETDuet-$LHGFR_{2N4C}$ | pETDuet-1 contained the gene of $LHGFR_{2N4C}$ |
| pETDuet-$LHGFR_{2N5C}$ | pETDuet-1 contained the gene of $LHGFR_{2N5C}$ |
| pETDuet-$LHGFR_{2N6C}$ | pETDuet-1 contained the gene of $LHGFR_{2N6C}$ |
| pETDuet-$LHGFR_{2N7C}$ | pETDuet-1 contained the gene of $LHGFR_{2N7C}$ |
| pETDuet-$LHGFR_{3N0C}$ | pETDuet-1 contained the gene of $LHGFR_{3N0C}$ |
| pETDuet-$LHGFR_{3N1C}$ | pETDuet-1 contained the gene of $LHGFR_{3N1C}$ |
| pETDuet-$LHGFR_{3N2C}$ | pETDuet-1 contained the gene of $LHGFR_{3N2C}$ |
| pETDuet-$LHGFR_{3N3C}$ | pETDuet-1 contained the gene of $LHGFR_{3N3C}$ |
| pETDuet-$LHGFR_{3N4C}$ | pETDuet-1 contained the gene of $LHGFR_{3N4C}$ |
| pETDuet-$LHGFR_{3N5C}$ | pETDuet-1 contained the gene of $LHGFR_{3N5C}$ |
| pETDuet-$LHGFR_{3N6C}$ | pETDuet-1 contained the gene of $LHGFR_{3N6C}$ |
| pETDuet-$LHGFR_{3N7C}$ | pETDuet-1 contained the gene of $LHGFR_{3N7C}$ |
| pETDuet-$LHGFR_{4N0C}$ | pETDuet-1 contained the gene of $LHGFR_{4N0C}$ |
| pETDuet-$LHGFR_{4N1C}$ | pETDuet-1 contained the gene of $LHGFR_{4N1C}$ |
| pETDuet-$LHGFR_{4N2C}$ | pETDuet-1 contained the gene of $LHGFR_{4N2C}$ |
| pETDuet-$LHGFR_{4N3C}$ | pETDuet-1 contained the gene of $LHGFR_{4N3C}$ |
| pETDuet-$LHGFR_{4N4C}$ | pETDuet-1 contained the gene of $LHGFR_{4N4C}$ |
| pETDuet-$LHGFR_{4N5C}$ | pETDuet-1 contained the gene of $LHGFR_{4N5C}$ |
| pETDuet-$LHGFR_{4N6C}$ | pETDuet-1 contained the gene of $LHGFR_{4N6C}$ |
| pETDuet-$LHGFR_{4N7C}$ | pETDuet-1 contained the gene of $LHGFR_{4N7C}$ |

| | |
|---|---|
| pETDuet-LHGFR$_{\text{G4-0N0C}}$ | pETDuet-1 contained the gene of LHGFR$_{\text{G4-0N0C}}$ |
| pETDuet-LHGFR$_{\text{0N0C-G4}}$ | pETDuet-1 contained the gene of LHGFR$_{\text{0N0C-G4}}$ |
| pETDuet-LHGFR$_{\text{G4-0N0C-G4}}$ | pETDuet-1 contained the gene of LHGFR$_{\text{G4-0N0C-G4}}$ |
| pETDuet-LHGFR$_{\text{KL-0N0C}}$ | pETDuet-1 contained the gene of LHGFR$_{\text{KL-0N0C}}$ |
| pETDuet-LHGFR$_{\text{0N0C-KL}}$ | pETDuet-1 contained the gene of LHGFR$_{\text{0N0C-KL}}$ |
| pETDuet-LHGFR$_{\text{KL-0N0C-KL}}$ | pETDuet-1 contained the gene of LHGFR$_{\text{KL-0N0C-KL}}$ |
| pETDuet-LHGFR$_{\text{G4-0N3C}}$ | pETDuet-1 contained the gene of LHGFR$_{\text{G4-0N3C}}$ |
| pETDuet-LHGFR$_{\text{0N3C-G4}}$ | pETDuet-1 contained the gene of LHGFR$_{\text{0N3C-G4}}$ |
| pETDuet-LHGFR$_{\text{G4-0N3C-G4}}$ | pETDuet-1 contained the gene of LHGFR$_{\text{G4-0N3C-G4}}$ |
| pETDuet-LHGFR$_{\text{KL-0N3C}}$ | pETDuet-1 contained the gene of LHGFR$_{\text{KL-0N3C}}$ |
| pETDuet-LHGFR$_{\text{0N3C-KL}}$ | pETDuet-1 contained the gene of LHGFR$_{\text{0N3C-KL}}$ |
| pETDuet-LHGFR$_{\text{KL-0N3C-KL}}$ | pETDuet-1 contained the gene of LHGFR$_{\text{KL-0N3C-KL}}$ |
| pETDuet-LHGFR$_{\text{G4-0N7C}}$ | pETDuet-1 contained the gene of LHGFR$_{\text{G4-0N7C}}$ |
| pETDuet-LHGFR$_{\text{0N7C-G4}}$ | pETDuet-1 contained the gene of LHGFR$_{\text{0N7C-G4}}$ |
| pETDuet-LHGFR$_{\text{G4-0N7C-G4}}$ | pETDuet-1 contained the gene of LHGFR$_{\text{G4-0N7C-G4}}$ |
| pETDuet-LHGFR$_{\text{KL-0N7C}}$ | pETDuet-1 contained the gene of LHGFR$_{\text{KL-0N7C}}$ |
| pETDuet-LHGFR$_{\text{0N7C-KL}}$ | pETDuet-1 contained the gene of LHGFR$_{\text{0N7C-KL}}$ |
| pETDuet-LHGFR$_{\text{KL-0N7C-KL}}$ | pETDuet-1 contained the gene of LHGFR$_{\text{KL-0N7C-KL}}$ |
| pcDNA3.1$^{(+)}$ | Vector for protein expression in human cells; Ap$^{r}$ |
| pcDNA3.1$^{(+)}$-LHGFR$_{\text{0N3C}}$ | pcDNA3.1$^{(+)}$ contained kozak sequence and the gene of LHGFR$_{\text{0N3C}}$ |
| pcDNA3.1$^{(+)}$-LHGFR$_{\text{0N7C}}$ | pcDNA3.1$^{(+)}$ contained kozak sequence and the gene of LHGFR$_{\text{0N7C}}$ |
| pTKRED | Plasmid expressing λRed recombinase genes for gene knock out; Spe$^{r}$ |

| | |
|---|---|
| pCP20 | Plasmid expressing Flp recombinase to remove kanamycin resistance cassette during gene knock out; $Cm^r$ |
| pKD4 | Template for amplification of the kanamycin resistance cassette; $Km^r$ |
| pEASY-Blunt | Vector for blunt-end cloning; $Ap^r$ |
| pEASY-Blunt-F1 | pEASY-Blunt contained F1 fragment |
| pEASY-Blunt-F2 | pEASY-Blunt contained F2 fragment |

[a] $G_4$ and KL indicate a flexible linker Gly-Gly-Gly-Gly-Ser-Gly-Gly and a rigid linker Lys-Leu-Tyr-Pro-Tyr-Asp-Val-Pro-Asp-Tyr-Ala, respectively, and they are added at the N-terminus or C-terminus of LhgR and its variants.

[b] $Tc^r$, tetracycline resistant; $Km^r$, kanamycin resistant; $Ap^r$, ampicillin resistant; $Spe^r$, spectinomycin resistant; $Cm^r$, chloramphenicol resistant.

**Supplementary Table 4 Oligonucleotides used in this study**

| Primer | Sequence(5'-3')[a] | Use[b] |
|---|---|---|
| **Expression** | | |
| *lhgO*-F | AATTGAATTCATGACATACGACTACTGCA (EcoRI) | Amplification of *lhgO* in *P. putida* W619 (forward) |
| *lhgO*-R | AATTGGTACCTCAGCTGGCCTTGAGGAT (KpnI) | Amplification of *lhgO* in *P. putida* W619 (reverse) |
| F2-*lhgR*-F1-*lhgO*-F | AATTGAGCTCAAGTCTGCCTGACGCGGCC (SacI) | Amplification of F2-*lhgR*-F1-*lhgO* in *P. putida* W619 (forward) |
| F2-*lhgR*-F1-*lhgO*-R | AATTGGATCCTCAGCTGGCCTTGAGGATCT (BamHI) | Amplification of F2-*lhgR*-F1-*lhgO* in *P. putida* W619 (reverse) |
| *lhgR*-F | AATTGGATCCGATGCTAGAACTCCAGC (BamHI) | Amplification of *lhgR* in *P. putida* W619 (forward) |
| *lhgR*-R | AATTAAGCTTTCAGTCGAGTGCAGGT (HindIII) | Amplification of *lhgR* in *P. putida* W619 (reverse) |
| mTFP-F | AATTGGATCCGATGGTGAGCAAGGGCGAGGAGA (BamHI) | Amplification of mTFP (forward) |
| mTFP-R | AATTGAGCTCCTTGTACAGCTCGTCCATGCCGT (SacI) | Amplification of mTFP (reverse) |
| Venus-F | AATTGTCGACATGGTGAGTAAAGGCGAAGAACTGT (SalI) | Amplification of Venus (forward) |
| Venus-R | AATTGCGGCCGCTTATTTATACAGTTCATCCATGCCC (NotI) | Amplification of Venus (reverse) |
| *lhgR*-0N | CGAGCTGTACAAGGAGCTCATGCTAGAACTCCAGCGCCC | Amplification of LhgR$_{0N}$ (forward) |
| *lhgR*-1N | CGAGCTGTACAAGGAGCTCCTAGAACTCCAGCGCCCCGA | Amplification of LhgR$_{1N}$ (forward) |
| *lhgR*-2N | CGAGCTGTACAAGGAGCTCGAACTCCAGCGCCCCGACAC | Amplification of LhgR$_{2N}$ (forward) |
| *lhgR*-3N | CGAGCTGTACAAGGAGCTCCTCCAGCGCCCCGACACTCT | Amplification of LhgR$_{3N}$ (forward) |
| *lhgR*-4N | CGAGCTGTACAAGGAGCTCCAGCGCCCCGACACTCTGGT | Amplification of LhgR$_{4N}$ (forward) |
| *lhgR*-0C | CTTTACTCACCATGTCGACGTCGAGTGCAGGTAGTTCTA | Amplification of LhgR$_{0C}$ (reverse) |
| *lhgR*-1C | CTTTACTCACCATGTCGACGAGTGCAGGTAGTTCTATTT | Amplification of LhgR$_{1C}$ (reverse) |

| | | |
|---|---|---|
| *lhgR*-2C | CTTTACTCACCATGTCGACTGCAGGTAGTTCTATTTTCA | Amplification of $LhgR_{2C}$ (reverse) |
| *lhgR*-3C | CTTTACTCACCATGTCGACAGGTAGTTCTATTTTCAGGC | Amplification of $LhgR_{3C}$ (reverse) |
| *lhgR*-4C | CTTTACTCACCATGTCGACTAGTTCTATTTTCAGGCGTT | Amplification of $LhgR_{4C}$ (reverse) |
| *lhgR*-5C | CTTTACTCACCATGTCGACTTCTATTTTCAGGCGTTTG | Amplification of $LhgR_{5C}$ (reverse) |
| *lhgR*-6C | CTTTACTCACCATGTCGACTATTTTCAGGCGTTTGGCAG | Amplification of $LhgR_{6C}$ (reverse) |
| *lhgR*-7C | CTTTACTCACCATGTCGACTTTCAGGCGTTTGGCAGAGG | Amplification of $LhgR_{7C}$ (reverse) |
| *lhgR*-8C | CTTTACTCACCATGTCGACCAGGCGTTTGGCAGAGGCG | Amplification of $LhgR_{8C}$ (reverse) |
| *lhgR*-9C | CTTTACTCACCATGTCGACGCGTTTGGCAGAGGCGCGCA | Amplification of $LhgR_{9C}$ (reverse) |
| *lhgR*-10C | CTTTACTCACCATGTCGACTTTGGCAGAGGCGCGCAGA | Amplification of $LhgR_{10C}$ (reverse) |
| *lhgR*-11C | CTTTACTCACCATGTCGACGGCAGAGGCGCGCAGATGCGCT TCG | Amplification of $LhgR_{11C}$ (reverse) |
| *lhgR*-12C | CTTTACTCACCATGTCGACAGAGGCGCGCAGATGCGCT | Amplification of $LhgR_{12C}$ (reverse) |
| *lhgR*-13C | CTTTACTCACCATGTCGACGGCGCGCAGATGCGCTTCG | Amplification of $LhgR_{13C}$ (reverse) |
| *lhgR*-14C | CTTTACTCACCATGTCGACGCGCAGATGCGCTTCGGCAC | Amplification of $LhgR_{14C}$ (reverse) |
| *lhgR*-15C | CTTTACTCACCATGTCGACCAGATGCGCTTCGGCACAC | Amplification of $LhgR_{15C}$ (reverse) |
| $G_4$-*lhgR*-0N | CGAGCTGTACAAGGAGCTCGGCGGAGGCGGAAGCGGCGG AATGCTAGAACTCCAGCGCCC | Amplification of $G_4SG_2$-$LhgR_{0N}$ (forward) |
| KL-*lhgR*-0N | CGAGCTGTACAAGGAGCTCAAGCTGTATCCTTATGATGTTC CTGATTATGCAATGCTAGAACTCCAGCGCCC | Amplification of KLYPYDVPDYA-$LhgR_{0N}$ (forward) |
| *lhgR*-0C-$G_4$ | CTTTACTCACCATGAGCTCTCCGCCGCTTCCGCCTCCGCCGT CGAGTGCAGGTAGTTCTA | Amplification of $LhgR_{0C}$-$G_4SG_2$ (reverse) |
| *lhgR*-0C-KL | CTTTACTCACCATGAGCTCTGCATAATCAGGAACATCATAA GGATACAGCTTGTCGAGTGCAGGTAGTTCTAT | Amplification of $LhgR_{0C}$-KLYPYDVPDYA (reverse) |

| | | |
|---|---|---|
| *lhgR*-3C-G$_4$ | CTTTACTCACCATGAGCTCTCCGCCGCTTCCGCCTCCGCCA GGTAGTTCTATTTTCAGGC | Amplification of LhgR$_{3C}$-G$_4$SG$_2$ (reverse) |
| *lhgR*-3C-KL | CTTTACTCACCATGAGCTCTGCATAATCAGGAACATCATAA GGATACAGCTTAGGTAGTTCTATTTTCAGGC | Amplification of LhgR$_{3C}$-KLYPYDVPDYA (reverse) |
| *lhgR*-7C-G$_4$ | CTTTACTCACCATGAGCTCTCCGCCGCTTCCGCCTCCGCCTT TCAGGCGTTTGGCAGAGG | Amplification of LhgR$_{7C}$-G$_4$SG$_2$ (reverse) |
| *lhgR*-7C-KL | CTTTACTCACCATGAGCTCTGCATAATCAGGAACATCATAA GGATACAGCTTTTTCAGGCGTTTGGCAGAGG | Amplification of LhgR$_{7C}$-KLYPYDVPDYA (reverse) |
| **Gene knockout in *P. putida* KT2440** | | |
| *lhgO*-uf | ATTGAATTCGGTGTACGATTTCATCATCATT (EcoRI) | Amplification of upstream homologous arm of *lhgO* (forward) |
| *lhgO*-ur | CAGGTCGCTGGGGCTACCGACGATGTTGGGCT | Amplification of upstream homologous arm of *lhgO* (reverse) |
| *lhgO*-df | CAACATCGTCGGTAGCCCCAGCGACCTGTTCC | Amplification of downstream homologous arm of *lhgO* (forward) |
| *lhgO*-dr | AATAAGCTTTCAGCTGGCAGCCCTCTGGTTG (HindIII) | Amplification of downstream homologous arm of *lhgO* (reverse) |
| *csiR*-uf | ATAGGATCCGATCTGATTGTGCCAAAGAA (BamHI) | Amplification of upstream homologous arm of *csiR* (forward) |
| *csiR*-ur | TGTGGGCCCGCTCTCCAAGCACCTG | Amplification of upstream homologous arm of *csiR* (reverse) |
| *csiR*-df | CAGGTGCTTGGAGAGCGGGCCCACA | Amplification of downstream homologous arm of *csiR* (forward) |
| *csiR*-dr | ATAGAATTCCATGACCTGGATCAGTGCG (EcoRI) | Amplification of downstream homologous arm of *csiR* (reverse) |
| *csiD*-uf | ATAGAATTCGATGAACGCCTTTACGCAGATCGAC (EcoRI) | Amplification of upstream homologous arm of *csiD* (forward) |

| | | |
|---|---|---|
| *csiD*-ur | CGAATACCGAATGGGATCGGGGTAATCAGCAT | Amplification of upstream homologous arm of *csiD* (reverse) |
| *csiD*-df | TGATTACCCCGATCCCATTCGGTATTCGACAC | Amplification of downstream homologous arm of *csiD* (forward) |
| *csiD*-dr | AATAAGCTTTTATTGACCGCGCTGGTACAGCGGC (HindIII) | Amplification of downstream homologous arm of *csiD* (reverse) |
| **Gene knockout in *E. coli* MG1655(DE3)** | | |
| *csiD*-F1 | CAGGAGAACCGCCGAACTAT | Amplification of upstream homologous arm of *csiD* (forward) |
| *csiD*-R1 | ACTTCGAAGCAGCTCCAGCCTACACCATCAGAAGCGATCCTCTTATGA | Amplification of upstream homologous arm of *csiD* (reverse) |
| *csiD*-F2 | TCATAAGAGGATCGCTTCTGATGGTGTAGGCTGGAGCTGCTTCGAAGT | Amplification of kanamycin resistance cassette for replacing *csiD* (forward) |
| *csiD*-R2 | CTTTGCGCTTACTGATGCGTCTGATGGGAATTAGCCATGGTC | Amplification of kanamycin resistance cassette for replacing *csiD* (reverse) |
| *csiD*-F3 | GACCATGGCTAATTCCCATCAGACGCATCAGTAAGCGCAAAG | Amplification of downstream homologous arm of *csiD* (forward) |
| *csiD*-R3 | CATTTCGAGATCGGACGTGG | Amplification of downstream homologous arm of *csiD* (reverse) |
| *lhgO*-F1 | CCGAGTAAAAACGTCAGCAAAGATG | Amplification of upstream homologous arm of *lhgO* (forward) |
| *lhgO*-R1 | ACTTCGAAGCAGCTCCAGCCTACACCATCCGCTCAATTCCTTTGCGCT | Amplification of upstream homologous arm of *lhgO* (reverse) |
| *lhgO*-F2 | AGCGCAAAGGAATTGAGCGGATGGTGTAGGCTGGAGCTGCTTCGAAGT | Amplification of kanamycin resistance cassette for replacing *lhgO* (forward) |
| *lhgO*-R2 | AGGTTATTGATTAAATGCGGCGTGATGGGAATTAGCCATGGTC | Amplification of kanamycin resistance cassette for replacing *lhgO* (reverse) |

| | | |
|---|---|---|
| *lhgO*-F3 | GACCATGGCTAATTCCCATCACGCCGCATTTAATCAATAACCT | Amplification of downstream homologous arm of *lhgO* (forward) |
| *lhgO*-R3 | TTTACCCTGTTCGAGGGTCATCAGG | Amplification of downstream homologous arm of *lhgO* (reverse) |
| **EMSAs** | | |
| F1-F | TACCCAGAGCTTGCTGCGAC | Amplification of DNA fragment F1 (forward) |
| F1-R | GCAGGGGTACCTTGTGATTCTT | Amplification of DNA fragment F1 (reverse) |
| F2-F | AAGTCTGCCTGACGCGG | Amplification of DNA fragment F2 (forward) |
| F2-R | GGAAGCGATTGCCTAGTGG | Amplification of DNA fragment F2 (reverse) |

[a]Restriction sites are underlined, and the restriction enzymes are indicated in parentheses.

[b]The subscript indicates the number of amino acids truncated from the N- or C-terminus of LhgR.
